# Single Cell Spatial Profiling Identifies Region-Specific Extracellular Matrix Adhesion and Signaling Networks in Glioblastoma

**DOI:** 10.1101/2024.10.31.621280

**Authors:** Arpan De, Santiago A. Forero, Ali Pirani, John E. Morales, Marisol De La Fuente-Granada, Sumod Sebastian, Jason T. Huse, Leomar Y. Ballester, Jeffrey S. Weinberg, Frederick F. Lang, Kadir C. Akdemir, Joseph H. McCarty

## Abstract

The human brain contains a rich milieu of extracellular matrix (ECM) components that are often dysregulated in pathologies including the malignant cancer glioblastoma (GBM). Here, we have used *in situ* single-cell spatial transcriptomic platforms to map the expression patterns of nearly 400 ECM genes in normal brain and GBM samples. Our analysis identifies at least four different GBM cell populations with unique ECM expression profiles that show spatial enrichment in distinct intratumor regions. Spatial mapping also demonstrates largely non-overlapping expression signatures of various ECM components in GBM stromal cell types, particularly in vascular endothelial cells and reactive microglia/macrophages. Comparisons of GBM (IDH1 wild type) versus lower-grade II and III astrocytoma samples (IDH1 R132H) identifies differential expression of key ECM components, including elevated levels of select ECM glycoproteins (IGFBP2 and MGP) and ECM-affiliated proteins (ANXA1 and ANXA2). In addition, we detect spatially enriched expression of COL8A1 (collagen), LUM (proteoglycan), and POSTN (ECM glycoprotein) in perivascular stromal cells in GBM but not in lower grade tumors. Computational analysis of putative ligand-receptor interactions reveals novel ECM communication networks between cancer cells and stromal components, particularly in regions of GBM microvascular proliferation and pseudopalisading necrosis. In summary, this comprehensive spatial map provides new insights into microenvironmental control of GBM initiation and progression and identifies potential therapeutic targets in the ECM.

## Introduction

Glioblastoma (GBM) is a primary brain cancer with rapidly proliferative and highly invasive growth features (1,2). Despite current treatments including surgery and chemoradiation, GBM typically recurs, resulting in a survival rate of about 12 to 15 months for most patients (3). Stromal cells in the GBM microenvironment, including reactive astrocytes, microglia and macrophages, and vascular endothelial cells, play key roles in promoting tumor growth and recurrence (4,5). Prior reports using single cell RNA-seq, proteomics, and/or spatial transcriptomic approaches have profiled the diverse stromal cell populations in GBM (6,7). GBM cells and their stromal counterparts also produce hundreds of extracellular matrix (ECM) factors that are a major component of the tumor stroma (8). ECM adhesion and signaling events are coupled to growth and progression of GBM; for example, stem-like cancer cells localize to ECM-rich vascular niches where they likely exploit local cues for growth and survival benefits (9). As GBM cells proliferate and infiltrate the brain parenchyma they remodel ECM components in vascular basement membranes to locally activate growth factor and cytokine signaling events, leading to pathological angiogenesis and edema due to the breakdown of the blood-brain barrier (BBB) (10). Invasive GBM cells also deposit new ECM factors in the interstitial matrices around neurons and perivascular spaces to form classic “secondary structures of Scherer” (11).

There are more than 1,000 genes that encode ECM proteins and related components that comprise the human matrisome (12). We understand surprisingly little about which matrisome genes are expressed in GBM and in lower grade brain tumors or how matrisome proteins facilitate contact and communication between transcriptionally diverse cancer cells and stromal cells. *In situ* spatial transcriptomic platforms, and in particular Xenium spatial profiling, can directly detect and quantify RNA transcripts within their native tissue environment (13).

Here, we have used Xenium-based single-cell spatial transcriptomics to profile the *in situ* expression of nearly 400 matrisome genes in human brain tumors, including grade II and III astrocytoma as well as multiple GBM samples. These efforts reveal a heterogeneous pattern of ECM gene expression, and show differential enrichment of many matrisome genes in cancer cells and stromal cell types in select spatial regions in GBM. Computational analyses identify several novel ECM ligand-receptor pairings that mediate adhesion and communication between cancer cells and stromal cells in the GBM microenvironment. Collectively, these efforts uncover new ECM factors in GBM as well as their specific cells of origin, which may lead to potential therapeutic interventions to improve survival of patients with GBM.

## Materials and Methods

### Human tissue samples

All human tissue samples were collected in accordance with the Institutional Review Board of The University of Texas MD Anderson Cancer Center, Houston, and subsequently de-identified for all experimental studies. Patient-derived samples included grade II glioma (n=9), grade III astrocytoma (n=11), GBM (n=8), normal (n=4), and tissue matched GBM core and normal (n=3) tissue samples.

### Xenium gene panel design and analysis

The panel was designed to detect mRNA expression of genes categorized as core matrisome and matrisome-affiliated (https://matrisomedb.org), in addition to cellular markers relevant to normal brain functions and brain tumor growth/progression. A total of 478 genes were selected and curated (Supplemental File #1) primarily based on single-cell atlas data ^43^ and the MatrisomeDB (https://matrisomedb.org). Fluorophore-tagged probes were designed by 10x Genomics (Pleasanton, CA). All parameters for Xenium analysis are manufacturer-designed and optimized ^16^.

### Cell segmentation

Grade III astrocytoma (n=1), GBM (n=1), and tissue matched normal and GBM core samples (n=3) were used for cell segmentation on the Xenium analyzer with Xeniumranger version 1.7.1.0 (10x Genomics, Pleasanton, CA) which utilizes nuclear expansion distance of 15 µm to establish cell boundaries. Grade II glioma (n=9), grade III astrocytoma (n=10), GBM (n=7) and non-cancerous brain tissues (n=4) were used for cell segmentation on the Xenium analyzer with Xeniumranger version 2.0.0.10 (10x Genomics, Pleasanton, CA) which utilizes nuclear expansion distance of 5 µm to establish cell boundaries.

### Downstream analysis

Single cell IDs for regions of interest were exported from the Xenium Explorer software (10x Genomics, Pleasanton, CA) as CSV files and used for downstream analysis. Off-instrument downstream analyses were performed on RStudio 4.4.0. Molecular coordinates and h5 matrix files from each xenium experiment were imported using the Seurat and scCustomize packages, and CSV files containing cell IDs for regions of interest were imported into R using the csv package (Bergsma 2022). Cluster analyses and differential gene expression were performed using built in functions from the Seurat package ^44^. Pseudo-bulk differential gene expression was performed using the DESeq2 package ^45^. Data integration was performed using the Harmony package (14).

### Spatial data Integration

Grade III astrocytoma and GBM data were integrated in R studio by merging the two data sets into one Seurat object and then performing the Harmony integration method which projects all cells into a shared embedding, which allows cells to be grouped by cell type rather than sample identity. Cluster analysis was performed by assessing the differential expression of conserved marker genes following the Seurat workflow for integrated single cell data.

### Annotating tumor regions of interest using Ivy GAP

Marker gene RNA-seq data from the IVY Glioblastoma Atlas Project (15) was used to identify anatomical features in the GBM sample. Using region-specific marker genes present in our Xenium panel, we identified Cellular Tumor (CT), Pseudopalisading (Pseudo.A, Pseudo.B), Microvascular proliferation (MVP), and Hyperplastic MVP regions.

### Cell-type and transcript abundance

The relative abundance of cell types was assessed by taking the proportion of each cell-type for a whole sample, or for a selected region of interest. Proportions were shown as voronoi plots, stacked barplots, or dotplots of percentages. Transcript abundance was assessed by taking the proportion of transcripts expressed in a region or cluster out of the total transcripts for a gene.

### Differential gene expression

Differential gene expression analyses in GBM, grade III astrocytoma, integrated grade III and GBM data, paired tumor vs. normal comparison, tumor grade comparison, and tissue matched grade III vs. normal (n=1), grade III vs. normal (n=2) and GBM vs. normal (n=1) were performed using the FindAllMarkers and FindMarkers functions in the Seurat workflow which identifies differentially expressed genes between groups of cells using a Wilcoxon Rank Sum test on normalized data counts. A log2fold change (log2FC) greater than 1 or less than -1 and an adjusted p-value (Bonferroni correction for multiple comparisons) of less than 0.05 were used as cutoffs for statistical significance. Differential gene expression analysis for patient brain tumor grade comparisons (grade II vs. GBM and grade III vs. GBM) was performed following the DESeq2 protocol for pseudo-bulk data (Harvard Chan Bioinformatics Core website).

### Z-score calculations

For grade III astrocytoma, the top 10 differentially expressed genes for glycoproteins, collagens, integrins, proteoglycans, and ECM-affiliated proteins marker genes were filtered for As-like, ECs, MG/MØs, OPC/OL-like, and Neu-like cells. Z-scores were calculated by taking the average normalized transcript count of a gene for each cell type or tumor subcluster minus the average normalized transcript count for all cell types or subclusters and dividing the difference over the standard deviation of all cell types or subclusters.

### CellChat cell-cell interactions

CellChat v2.1.0 was used to analyze region-specific cell-cell interactions and identify key signaling pathways along with their corresponding ligand-receptor pairs. The analysis was conducted in a contact- or spatial distance-dependent manner, utilizing the truncated mean method to compute average gene expression and communication probabilities of signaling pathways. Additionally, the default ‘CellChatDB.human’ database was customized to incorporate MGP pathways, designating MGP as the ligand and ITGB1, ITGAV, ITGB3, TGFB1, TGFBI, and BMPER as its receptors.

### Immunohistochemistry

Immunohistochemical analyses were performed with the following primary antibodies: rabbit anti-annexin A2 (1:200; catalog #11256–1-AP, Proteintech) and mouse anti-fibronectin antibody (1:300, catalog #66042-1-Ig, Proteintech). Slides were incubated in ABC Reagent (catalog #PK-4000, Vector Laboratories) for 30 min at RT. ImmPACT DAB Substrate (catalog #SK-4105, Vector Laboratories) was added for signal detection.

## Results

Single-cell spatial mRNA profiling was performed using multiple human astrocytoma sections of differing grades as well as matching non-cancerous human brain samples (Fig. 1A and Supp. Table 1). Custom-designed DNA probes recognizing 478 gene products encoding core ECM factors (n=148), ECM-affiliated factors (n=167), integrins (n=23), and other selected markers enriched in tumor and/or brain stromal cell types (n=140) were used to profile gene expression (Supp. Table 2). Unsupervised single-cell clustering by uniform manifold approximation and projection (UMAP) analyses identified multiple tumor and stromal cell types in GBM (Fig. 1B-D). Detailed analysis of the tumor cell (TC) population identified four distinct subclusters based on differential expression profiles of matrisome markers, tumor-enriched markers, and cell proliferation markers (Figs. 1E-F and 2B). Available RNA-seq data from the Ivy Glioblastoma Atlas Project (GAP) (15) were cross-referenced to compare gene expression profiles across four GBM histopathological regions to select gene signatures exhibiting pattern enrichment (Fig. 1G-H). Analysis of spatial single-cell transcriptional enrichment profiles in TC subtypes uncovered four anatomically and spatially distinct histopathologic regions within the GBM sample (Figs. 1G-H and 2A). TC subtypes 1 and 3 were more diffusely positioned throughout the tumor in regions annotated as cellular tumor (CT) and microvascular proliferation (MVP), respectively (Figs. 1H and 2A). In contrast, TC subtypes 2 and 4 were more enriched near pseudopalisading regions Pseudo.A and Pseudo.B, respectively.

**Figure 1.**
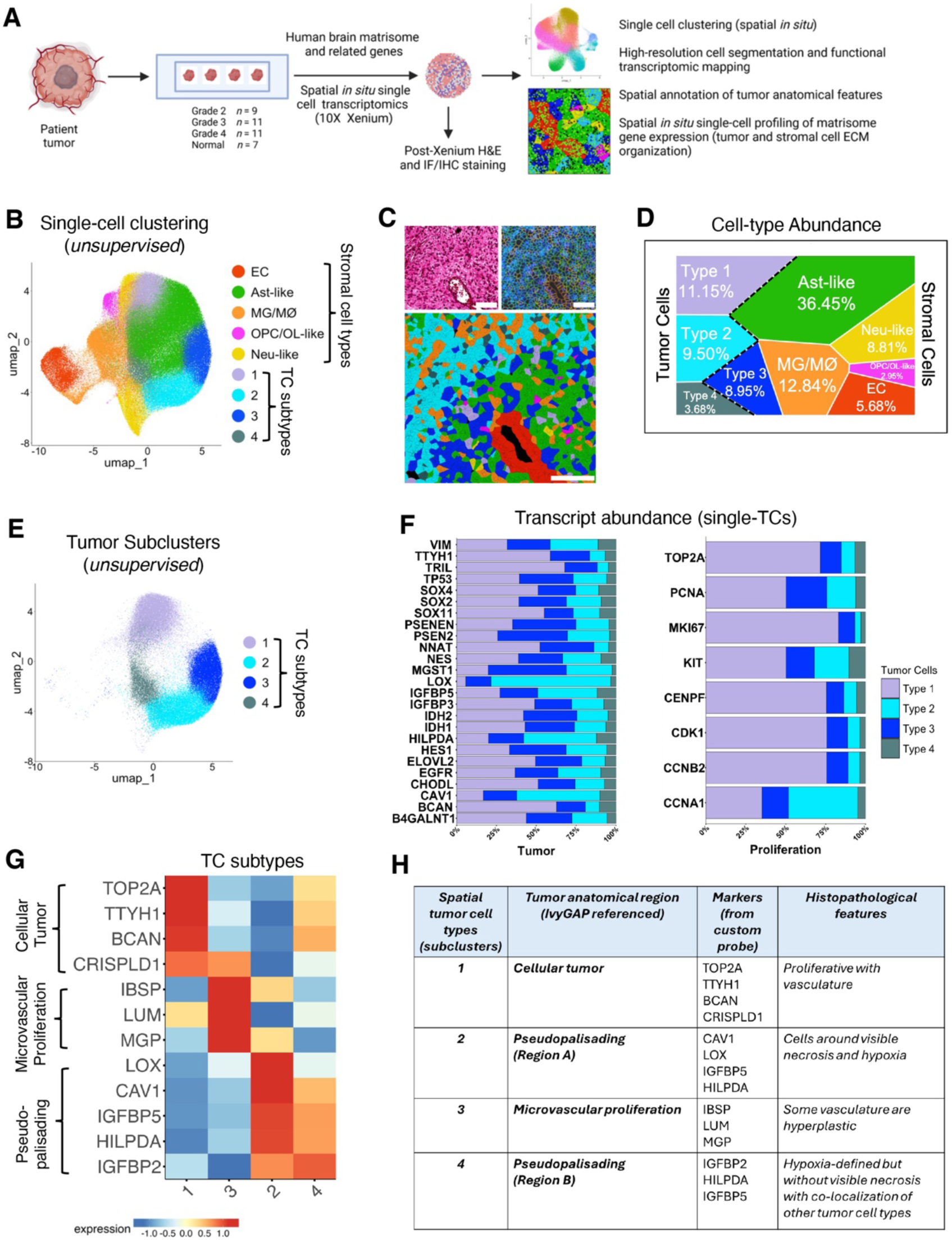
*In situ* single-cell analysis of matrisome gene expression in GBM. **(A);** Experimental design showing patient samples and strategies for in situ spatial profiling efforts. A total of 38 patient samples (differentially graded brain tumor and normal) were profiled. A custom probe panel was designed to detect expression of core matrisome, ECM-related, tumor-enriched, and cell-type markers (n=478) of the human brain. **(B);** Uniform manifold approximation and projection (UMAP) visualization representing unsupervised single-cell fine clustering of tumor and stromal cell types annotated from a human GBM sample. Pseudo-colored clusters are defined by differential expression profiles for cell-type specific and related matrisome markers. **(C);** Representative region from a GBM sample showing H&E stained (left, upper panel) and high-resolution cell segmentation images (right, upper panel). Bottom panel shows a magnified image of the region with single cell-types assigned pseudo-colors based on UMAP annotations (B). Scale bars: 100 µm. **(D);** Voronoi diagram showing abundances of tumor and stromal cell-type clusters constituting the GBM sample. Quantitation of relative percentages based on the total number of cells (n=321,020). **(E);** Subset UMAP showing the four TC subclusters. **(F);** Stacked bar plot showing single-cell transcript abundance for TC enriched genes (left) and cell proliferation markers (right) for the four subtypes identified in the GBM sample (E). **(G);** Expression heatmap showing top DEGs in TC subtypes used to annotate GBM anatomic features. Marker genes selected from the Ivy Glioblastoma Atlas Project (GAP) database. Normalized expression values were averaged by tumor subtype, log-transformed and scaled. **(H);** Summary of the histopathological features identified for each anatomic region in the GBM tissue annotated by the enrichment of select markers detected in the matrisome panel (G).

**Figure 2.**
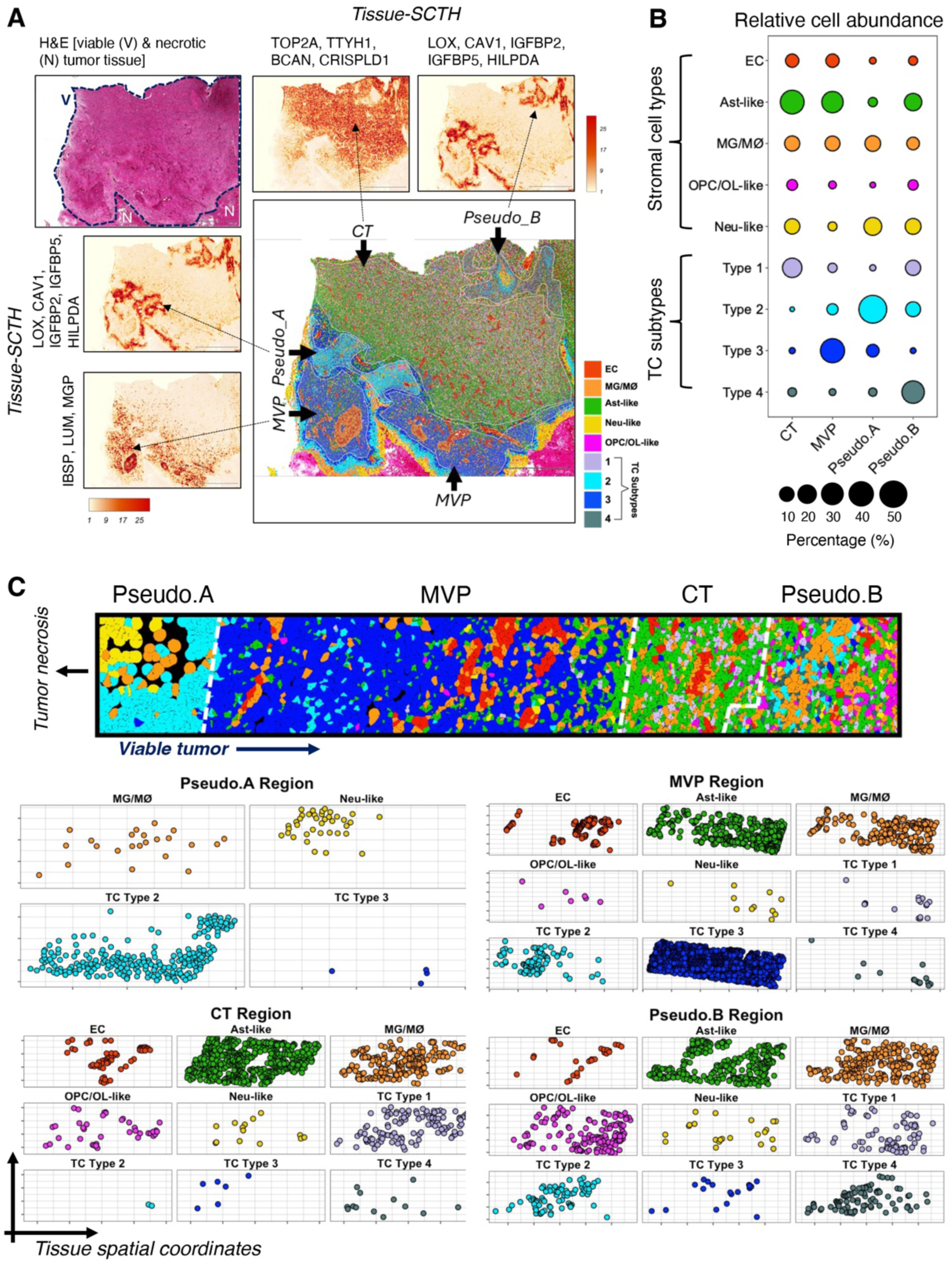
Spatial organization of TCs and stromal cell types in GBM. **(A);** Representative H&E-stained region of a GBM sample showing viable (V) and necrotic (N) tumor tissue (top left). Only viable tissue (encircled by dotted line) was analyzed. Spatial cumulative transcriptomic heatmap (SCTH) showing localization and co-enrichment of select markers (Fig. 1G, H) used for annotating anatomical regions in the GBM tissue. Cellular tumor (CT), microvascular proliferation (MVP), and pseudopalisading (Pseudo.A and Pseudo.B) regions are indicated in the image showing pseudo-colored TC and stromal cell types. Scale bars: 2000 µm. **(B);** Dot plot representing relative abundance of TC and stromal cell-types from each anatomical region profiled across a representative GBM tissue subsection. Dot size corresponds to the percentage of TC subtypes and stromal cell-types per region. Total number of cells profiled per region; CT: 37,192, MVP: 8,978, Pseudo.A: 4,632, and Pseudo.B: 13,893. **(C);** Representative localization of TC and stromal cell types highlighting spatial organization and abundance. Single-cells are denoted by cell-type specific pseudo-colored solid circles across tissue coordinates within segregated GBM anatomical niches. The regions (Pseudo.A, MVP, CT and Pseudo.B) are indicated by white dashed lines (top). Two-dimensional layered representation of spatial cellular organization in each GBM anatomical landscape (bottom). Cell coordinates were extracted using the 10X Genomics Xenium Explorer software.

We next analyzed the spatial distribution of TC and stromal cell types by sampling a tissue area comprising 64,695 single cells within a GBM sample, containing all four anatomic regions (Fig. 2B). The proportion of cell types and their corresponding *in situ* spatial organization showed distinctive patterns of localization (Fig. 2B-C).

To analyze tumor grade-specific differences in matrisome gene expression, we performed parallel spatial transcript mapping of a grade III astrocytoma sample (IDH1 R132H). Spatial transcript profiling and unsupervised single cell clustering by UMAP analysis identified diverse TC and stromal cell types in the grade III astrocytoma (Supp. Fig. 1A-B). UMAP analysis revealed the presence of four TC sub-clusters based on differential gene expression (Supp. Fig. 1A-C). Several differentially expressed genes (DEGs) linked to cell proliferation, e.g. MKI67 and PCNA, were enriched in TC clusters 1 and 4 in comparison to clusters 2 and 3 (Supp. Fig. 1D). Several DEGs belonging to different ECM categories of the matrisome were more enriched in TC cluster 4 (Supp. Fig. 1E). Analysis of spatial positioning of the four different TC clusters revealed largely uniform distribution across the grade III astrocytoma sample (Supp. Fig. 1F-G).

TCs and stromal cells in the four anatomic regions (CT, MVP, Pseudo.A, and Pseudo.B) of the GBM sample were next analyzed. Subtype 4 TCs populating the hypoxic Pseudo.B region revealed comparable gene expression profiles with subtype 1 TCs, localized in close spatial proximity to the CT region (Figs. 2 and 3A). Subtype 1 and 4 TCs showed comparable enrichment of ECM genes ELFN2, COL20A1, GPC2, TNR and CLEC7A (Fig. 3A). In contrast, transcripts for HAPLN1, COL9A1, COL9A3, FBN3 and CSPG5 were more abundant in subtype 1 TCs within the CT region. PLXNA4, PLXNB3, SEMA5B and CLEC5A transcripts displayed elevated expression in the Pseudo.B region comprising subtype 4 TCs (Fig. 3A). Similarly, subtype 2 TCs populating the Pseudo.B region were located near necrotic borders and within the hypoxic microenvironment of MVP region populated by subtype 3 TCs (Figs. 2A and 3A).

**Figure 3.**
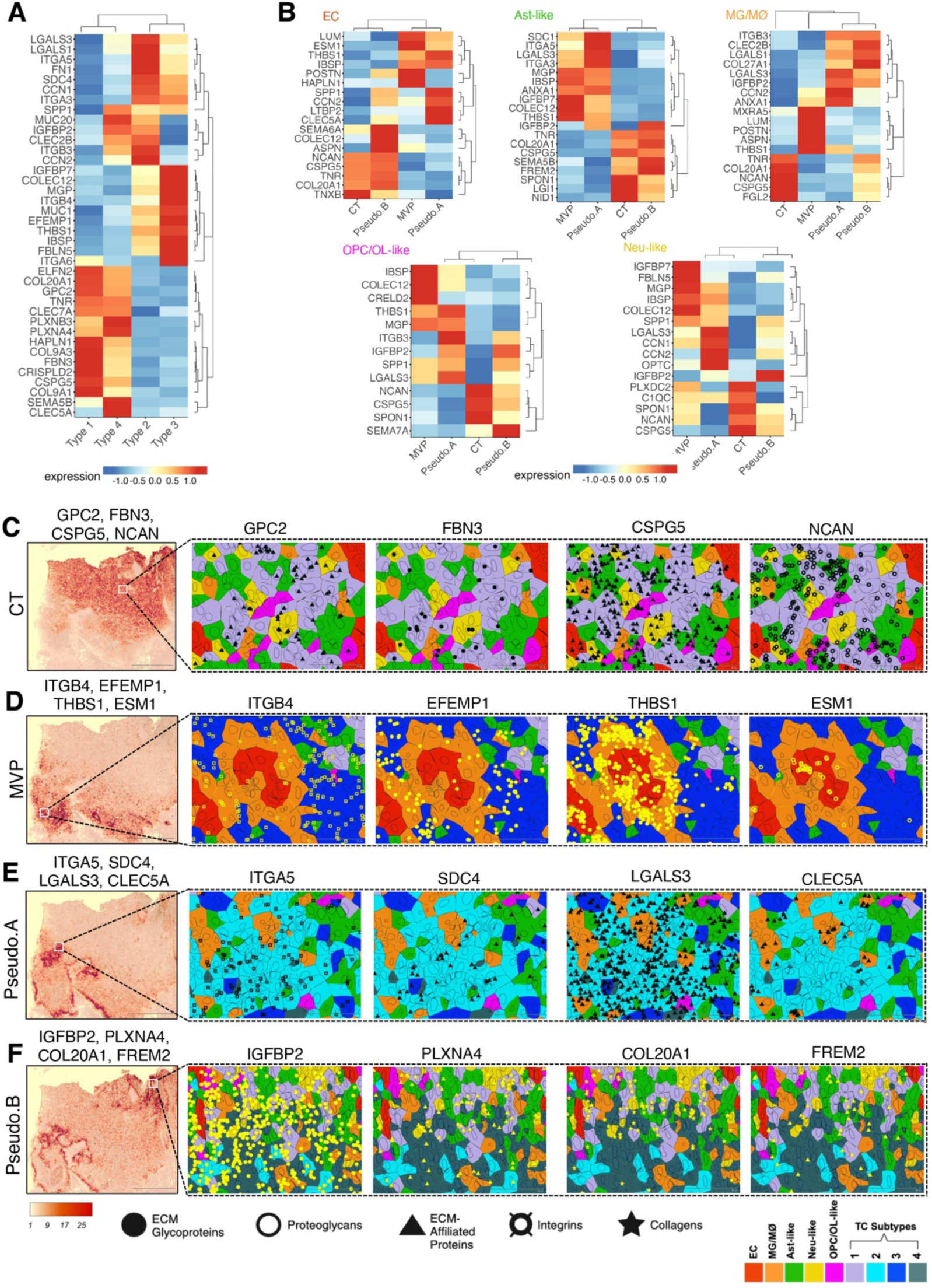
Matrisome gene expression profiles in TCs and stromal cells reveal region specificity. **(A);** Heatmap showing top DEGs in TC subtypes in different GBM regions. TC subtypes residing in different anatomic regions reveal enrichment of unique ECM signatures. Normalized expression values were averaged by tumor subtypes, log-transformed and scaled. **(B);** Expression heatmaps showing top DEGs in single stromal cells within the four annotated GBM regions (CT, MVP, Pseudo.A, Pseudo.B). Analysis was performed separately for each stromal cell type. Normalized expression values were averaged by region, log-transformed and scaled. **(C-F);** Spatial *in situ* transcript localization showing expression patterns of select ECM genes (from A and B) at single-cell resolution in various GBM regions. Magnified images reveal distinct localization of these ECM signatures in TC subtypes and nearby stromal cell types in regions of interest, shown as boxed areas within SCTHs for select gene groups indicated for CT (C), MVP (D), Pseudo.A (E) and Pseudo.B (F) landscapes in the GBM tissue. Transcript shape signifies gene group category (see legend in figure). Scale bars: 2000 µm and 50 µm.

While certain ECM transcripts, namely IGFBP7, COLEC12, MGP, ITGB4, THBS1, IBSP and FBLN5, showed higher expression in subtype 3 TCs in the MVP region, transcripts for LGALS1, LGALS3, ITGA5 and FN1 were abundant in Pseudo.B region containing subtype 2 TCs (Fig. 3A).

Comprehensive analysis of stromal populations identified microglia (MG), vascular endothelial cells (ECs), neural cells (Neu), oligodendrocyte progenitor cells (OPCs), oligodendrocytes (OLs), and astrocytes in both GBM and grade III astrocytoma samples (Figs. 1B-D and Supp. Fig. 1). Given that the higher grade astrocytomas are derived from transformed astroglia/neural stem cells of origin, we will refer to the non-cancerous tumor-infiltrating astrocyte population as “astrocyte-like” (Ast-like). This population likely contains a mixture of reactive non-malignant astrocytes and TCs that express astrocyte markers. Furthermore, the microglial population likely contains macrophages infiltrating from the circulation that express common markers and thus will be referred to as “microglia/macrophages” (MG/MØs). We detected diverse ECM gene expression profiles in the different GBM stromal cell types (Figs. 2 and 3B). ECs and Ast-like cells in both CT and Pseudo.B regions showed comparable expression profiles for some ECM genes (Fig. 3B). In contrast, ECs within the hypoxic Pseudo.B region showed enhanced expression for the ECM genes SEMA6A, COLEC12 and ASPN while IGFBP2, SEMA5B and FREM2 exhibited higher transcript localization in Ast-like cells (Fig. 3B). Similarly, ECs within the MVP region showed higher abundance of ECM genes LUM, ESM1, POSTN and HAPLN1 whereas those residing in the Pseudo.A region displayed transcript enrichment for SPP1, CCN2, LTBP2 and CLEC5A (Fig. 3B). The Pseudo.A region showed enrichment for SDC1, LGALS3, ITGA3 and ITGA5 transcripts. Tumor infiltrating MG/MØs reveal distinct ECM expression profiles in the CT and MVP regions (Fig. 3B). MG/MØs of the MVP region were enriched for some ECM genes namely LUM, POSTN and MXRA5, whereas TNR, COL20A1, CSPG5, NCAN and FGL2 transcripts displayed higher expression in MG/MØs in the CT region.

OPC/OL cells in the CT region showed increased expression of NCAN, CSPG5, and SPON1 transcripts whereas those in the MVP region had higher expression of IBSP, COLEC12 and CRELD2 (Fig. 3B). Higher transcript abundance of ITGB3 and LGALS3 were observed in OPC/OL cells in the Pseudo.A region in comparison to SEMA7A which displayed increased expression in Pseudo.B region. Some ECM genes were selectively enriched in Neu cells residing in the annotated GBM anatomical regions (Fig. 3B). TC subtypes and the stromal cell types revealed distinctly unique spatial expression patterns (Fig. 3C-F). For example, GPC2 and FBN3 were enriched in subtype 1 TCs in the CT region in comparison to CSPG5 and NCAN that revealed higher expression in the surrounding stromal cells in addition to the TCs (Fig. 3C). In the MVP region, ITGB4 and EFEMP1 transcripts were more abundant in subtype 3 TCs whereas ESM1 and THBS1 were more localized in proliferating microvascular ECs and perivascular MG/MØs (Fig. 3D). In the hypoxic Pseudo.A region, ITGA5 and SDC4 showed enhanced abundance in subtype 2 TCs in comparison to CLEC5A which was enriched in tumor infiltrating MG/MØs, and LGALS3 that was overall higher in both TCs and surrounding stromal cell types (Fig. 3E). In the Pseudo.B region, IGFBP2 transcripts were found in subtype 4 TCs but other ECM genes like PLXNA4, COL20A1 and FREM2 were elevated in stromal cell types surrounding the necrotic areas (Fig. 3F).

We next compared DEGs in the various stromal cell types in grade III astrocytoma versus GBM. The ten most DEGs encoding ECM glycoproteins were enriched in vascular ECs and in MG/MØs whereas overall expression levels of collagen genes were higher in Neu and OPC/OL cells (Supp. Fig. 2A). While the top ten differentially expressed integrin and proteoglycan genes were distributed between multiple cell types, ECM-affiliated genes showed highest enrichment in ECs followed by MG/MØs and Neu cells (Supp. Fig. 2A). There were several transcripts showing elevated expression in GBM stromal cell types versus stromal cell types in the grade III tumor, including IGFBP2 (ECM glycoprotein), ANXA1 (ECM-affiliated), ITGA7 (integrin), and SRGN (proteoglycan). Interestingly, we detected upregulation of specific matrisome factors such as MMRN1, SEMA7A, ITGA2, and ESM1 uniquely in GBM vascular ECs (Supp. Fig. 2B, C).

UMAP plots confirmed upregulation of select ECM glycoprotein genes, including IGFBP2 and MGP, in the GBM sample versus grade III astrocytoma (Supp. Fig. 3B-C). Spatial analysis of IGFBP2 and MGP transcript expression revealed upregulated expression in TCs, ECs, and Ast-like populations surrounding blood vessels in GBM (Supp. Fig. 3C). In comparison to vascular EC-specific expression in the grade III tumor, there was markedly elevated FN1 mRNA abundance in ECs and perivascular cells in the GBM sample (Supp. Fig. 3D-F). The integrin genes ITGA5 and ITGA7 showed significantly enhanced transcript counts in the GBM tissue compared to the grade III tumor sample (Supp. Fig. 4A-B). For example, elevated spatial expressions of ITGA5 transcripts were detected primarily in tumor ECs whereas ITGA7 transcripts were expressed mainly in TCs and Ast-like cells (Supp. Fig. 4B).

Among the 19 collagen gene transcripts that were profiled, COL16A1, COL8A1 and COL2A1 exhibited higher levels in the GBM tissue versus the grade III astrocytoma sample (Supp. Fig. 4C-D). There was increased spatial distribution of COL16A1 transcripts primarily in vascular ECs and MG/MØs, with lower expression in surrounding TCs and Ast-like populations in the GBM sample (Supp. Fig. 4D). Similarly, unlike in grade III astrocytoma, COL8A1 transcripts showed enhanced expression primarily in perivascular MG/MØs in GBM. We also detected increased COL8A1 transcript expression in GBM versus grade III astrocytoma based on UMAP analysis (Supp. Fig. 4D). Quantitative profiling of 25 genes encoding proteoglycans revealed that specific transcripts, including LUM and SRGN, exhibit significantly higher counts in the viable region of the GBM sample as determined by UMAP analyses (Supp. Fig. 4E-F). We detected enhanced spatial distribution of LUM primarily in GBM ECs and SRGN transcripts mainly in MG/MØs (Supp. Fig. 4F).

The ECM-affiliated genes ANXA1 and ANXA2 showed robust spatial enrichment in GBM compared to the grade III tumor (Supp. Fig. 4G-H). There was markedly enhanced expression of ANXA1 and ANXA2 transcripts primarily in GBM cells residing in close juxtaposition with vascular ECs (Supp. Fig. 4H). In comparison to the grade III sample, ECM affiliated genes (COLEC12, SDC2 and SEMA5A) showed enhanced spatial expression in GBM regions with perivascular microgliosis associated with pseudopalisading necrosis, that are largely absent in lower grade gliomas (Supp. Fig. 5A-C). Additionally, in comparison to the surrounding Ast-like cell populations, the ECM-affiliated gene LGALS3 revealed significantly elevated transcript localization in the pseudopalisading TCs near necrosis (Supp. Fig. 5D-E). As expected, these four ECM-affiliated genes (COLEC12, SDC2, SEMA5A and LGALS3) did not reveal obvious changes in spatial distribution of cellular transcripts in the grade III astrocytoma sample (Supp. Fig. 5F). We detected lower levels of transcripts for some matrisome genes in GBM versus grade III astrocytoma (Supp. Fig. 5G-J).

Next, histologically enriched markers from the Ivy-GAP dataset were used to annotate “hyperplastic” subregions within the GBM MVP area (Fig. 4A). These subregions contained perivascular MG/MØs showing enrichment for POSTN, COL8A1 and OGN transcripts in comparison to the adjacent non-hyperplastic vasculature within the MVP region (indicated by arrows in Fig. 4A). Single-cell transcript profiling for ECs and MG/MØs from multiple regions of interest for hyperplastic, MVP (non-hyperplastic) and CT regions revealed enrichment of diverse ECM genes (Fig. 4B). Several DEGs in ECs and MG/MØs were selected for further analysis.

**Figure 4.**
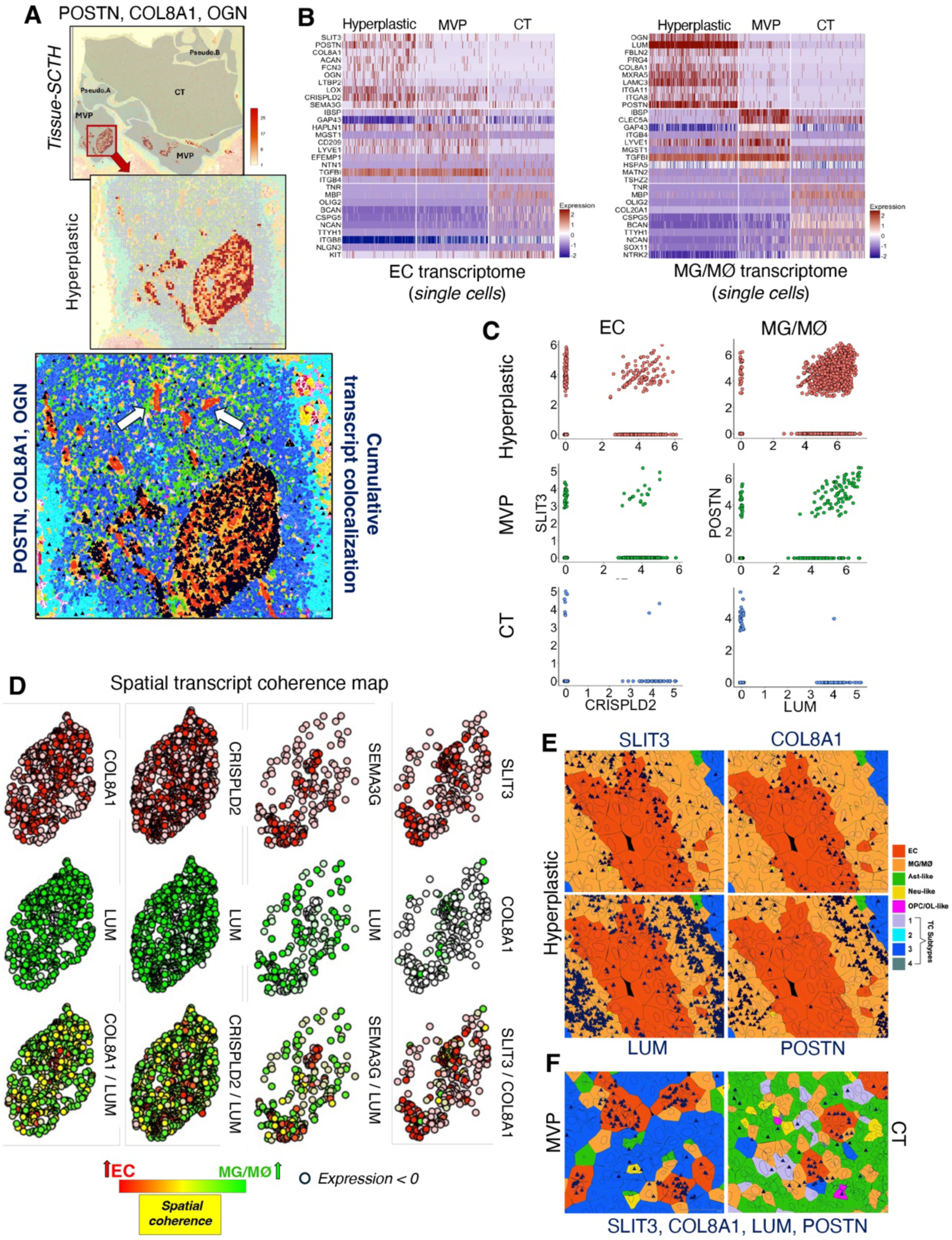
Heterogeneous matrisome gene expression in GBM blood vessels and perivascular MG/MØs. **(A);** SCTH of the GBM tissue showing spatial enrichment **\** in hyperplastic blood vessels in the MVP region (top). Magnified image (middle) shows a hyperplastic blood vessel with glomeruloid features in the boxed area (top). The bottom panel shows spatial co-localization of POSTN, COL8A1 and OGN transcripts (dark blue) in ECs and perivascular MG/MØs populating the hyperplastic blood vessels (white arrows). TC and stromal cell types are shown in pseudo-colors indicated in Figs. 1 and 2. Scale bars: 2000 µm and 500 µm. **(B);** Single-cell transcriptomic heatmaps showing top DEGs in ECs (left) and MG/MØs (right) residing in hyperplastic blood vessels (2,628 cells) as well as non-hyperplastic blood vessels in MVP (1,852 cells) and CT (2,226 cells) regions. **(C);** Feature scatter plots showing correlative expression of select genes from (B). **(D);** Representative coherence maps showing “spatial spots” for EC-expressed (red) and MG/MØ-expressed (green) transcripts for select genes revealing increased correlations in hyperplastic vascular regions in GBM. Enhanced spatial coherence is denoted by yellow “spatial spots” (bottom) for each gene set. **(E, F);** Spatial localization of transcripts (dark blue) for SLIT3, COL8A1, LUM and POSTN in ECs and MG/MØs localized in hyperplastic blood vessels (E), and MVP blood vessels located in the CT region (F). Scale bars: 50 µm.

ECM glycoproteins SLIT3 and CRSIPLD2 showed co-expression in ECs in hyperplastic blood vessels in comparison to ECs in the non-hyperplastic MVP and CT vasculature (Fig. 4C). The ECM-affiliated gene SEMA3G while COL8A1 in MG/MØs showed spatial coherence with SLIT3 enriched in hyperplastic ECs (Fig. 4D). In comparison to tumor vasculature in MVP and CT regions, SLIT3, COL8A1, LUM and POSTN revealed enhanced spatial enrichment within the GBM MVP region (Fig. 4E-F).

The CellChat tool (16) was used to study matrisome interaction networks via known ligand-receptor pairs and their associated signaling pathways (Fig. 5A-B). TC subtype 2 located in the Pseudo.A regions showed robust interactions with EC and MG/MØ populations localized in the MVP region primarily through SPP1 and FN1 signaling networks (Fig. 5B-C). The SPP1 network revealed strong outgoing and incoming signaling patterns from TC subtypes 2 and 3 in the Pseudo.A region to EC and MG/MØ populations residing in the adjacent MVP region, respectively. The FN1 network showed significant bi-directional communication patterns across these cell types localized in the adjacent GBM regions (Fig. 5C). The MVP-CT boundary in the GBM sample also revealed strong cellular interaction patterns (Fig. 5D-E). Ast-like and subtype 3 TCs communicated with adjacent MG/MØ populations across the MVP-CT boundary regions via the FN1 signaling network (Fig. 5E-F). Ast-like cell populations in CT and MVP regions strongly interacted through the laminin signaling network, which was also crucial in mediating cell-cell communication patterns between subtype 3 TCs in the MVP region, and closely localized Ast-like and subtype 1 TCs localized in the CT region (Fig. 5E-F). TCs in the CT region strongly communicated with closely juxtaposed perivascular Ast-like cells via collagens. Interestingly, TCs and stromal cells residing near the neighboring CT-Pseudo.B regions did not show robust cellular interactions (Supp. Fig. 7C-D).

**Figure 5.**
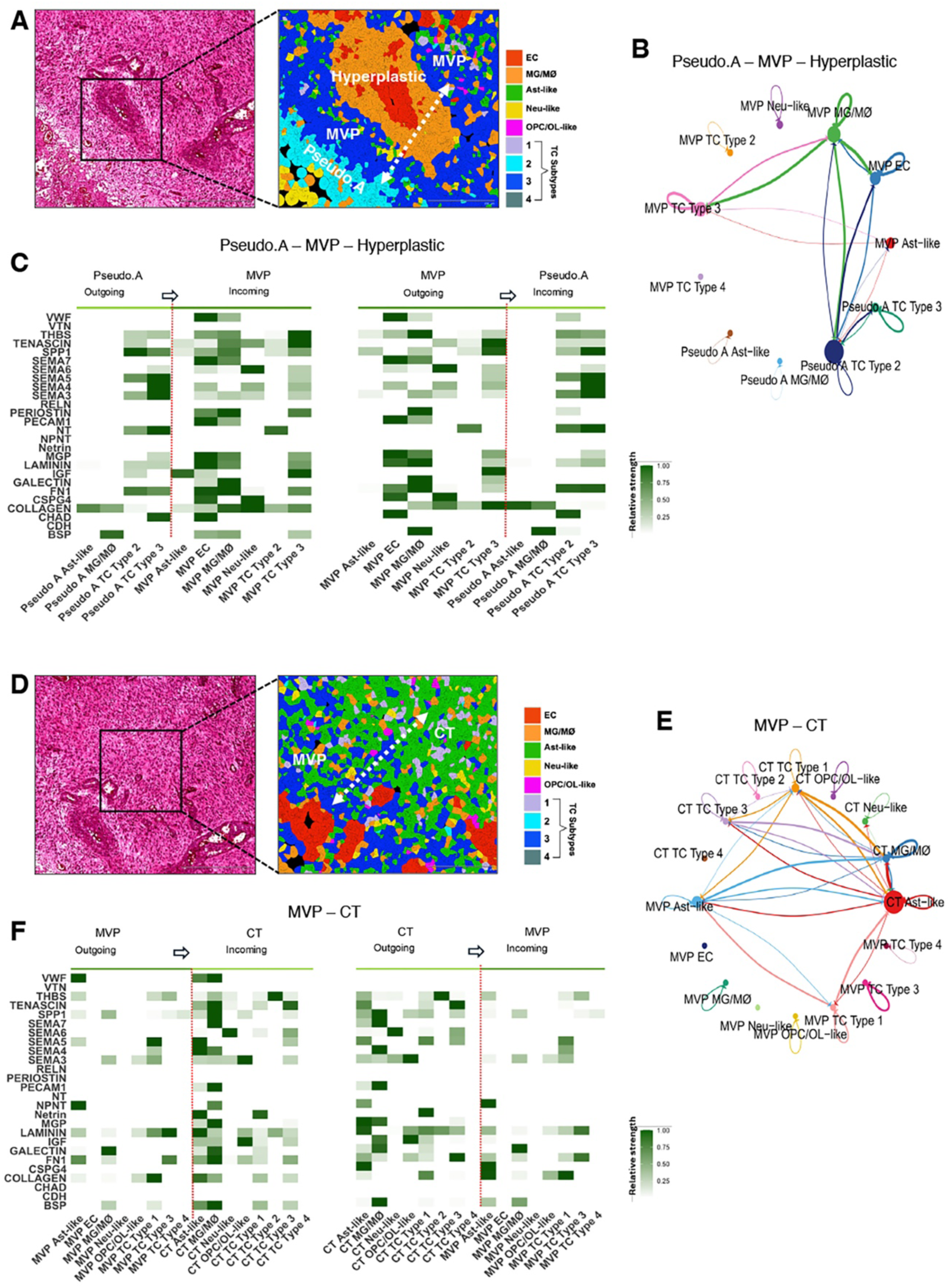
Cell-Cell communication networks across GBM anatomic regions. **(A**); H&E-stained image showing a Pseudo.A-MVP boundary including hyperplastic vasculature (boxed region) in the GBM sample (left). Magnified image (right) of the boxed area shown in the H&E-stained image (left) is high-resolution spatial annotation of GBM regions (Pseudo.A, MVP and Hyperplastic) and UMAP-based spatially segmented pseudo-colored (indicated) cell types. Dashed arrow signifies cellular crosstalk across “boundaries” spatially segregating the annotated GBM regions. Scale bars: 500 µm and 200 µm. **(B);** Chord plot summarizing interactions between specific cell types across Pseudo.A-MVP regions that include hyperplastic blood vessels. Each node (circle) represents a cell type, with its size equating to overall communication strength. Edges (lines) between nodes represent ligand-receptor interactions with the directionality of interaction indicated by arrows, pointing from the signaling (ligand-expressing) cell to the receiving (receptor-expressing) cell. Notably, MG/MØ and ECs within the MVP region show stronger interactions with sub-type 2 TCs in the Pseudo A region. **(C);** Heatmap of signaling networks showing outgoing and incoming signaling strength across the Pseudo A-MVP/Hyperplastic region. Color intensity indicates strength of ligand-receptor interactions within each pathway, stronger signals shown in darker green shades. Left panel shows outgoing signals from Pseudo A cell types (light green) to recipient MVP region cell types (dark green). The directionality of outgoing to incoming signaling interactions are displayed above each panel. Right panel shows outgoing signals from MVP region cell types (dark green) to recipient Pseudo A cell types (light green). **(D);** H&E-stained image showing MVP-CT boundary (boxed area) in the GBM tissue (left). Magnified image (right) of the boxed area shown in the H&E-stained image (left) represents high-resolution spatial annotation of GBM regions (MVP and CT) and UMAP-based pseudo-colored cell types. Dashed arrow signifies representative cellular crosstalk across “boundary” spatially segregating the annotated GBM niches indicated. Scale bars: 500 µm and 200 µm. (**E);** Chord plot summarizing interactions between region-specific cells across the MVP-CT boundaries. Excluding TC sub-types 2 and 4, cell types in the CT region engage in bidirectional crosstalk with varying strengths. **(F);** Heatmap of signaling networks showing outgoing and incoming signaling strength across MVP-CT region. Left panel shows outgoing signals from MVP cell types (dark green) to recipient CT region cell types (light green). Right panel shows outgoing signals from CT region cell types (light green) to recipient MVP cell types (light green).

Next, ligand-receptor pairs involved in intercellular signaling pathways were identified for “within” and “boundary” regions across different GBM regions (Fig. 6 and Supp. Fig. 6-7). The most notable ligand-receptor pairs were in the Pseudo.A-MVP-Hyperplastic and MVP-CT GBM anatomic regions (Fig. 6A, D). FN1 (ligand) enriched in/around proliferating ECs, and perivascular MG/MØs in the MVP region interacted with integrin receptors (ITGAV and ITGB8) on subtype 2 TCs localized in the neighboring Pseudo.A region (Fig. 6A-B). Bi-directional communication patterns were observed across the subtype 2 TCs (Pseudo.A region) and ECs (MVP region) via LAMB1 (ligand) and integrin receptors (ITGA7 and ITGB1) (Fig. 6A, C). Similar bi-directional interactions were detected across Ast-like cells (MVP region) and MG/MØ populations (CT region) through SPP1 (ligand) and corresponding integrin receptors (ITGAV and ITGB1) localized in both cell types (Fig. 6D-E). Additionally, Ast-like cells (MVP region) revealing enrichment of THBS4 (ligand) communicate with syndecan ligands (SDC1 and SDC4) localized in subtype 1 TCs and Ast-like cells respectively, residing in the neighboring GBM CT region (Fig. 6D, F).

**Figure 6.**
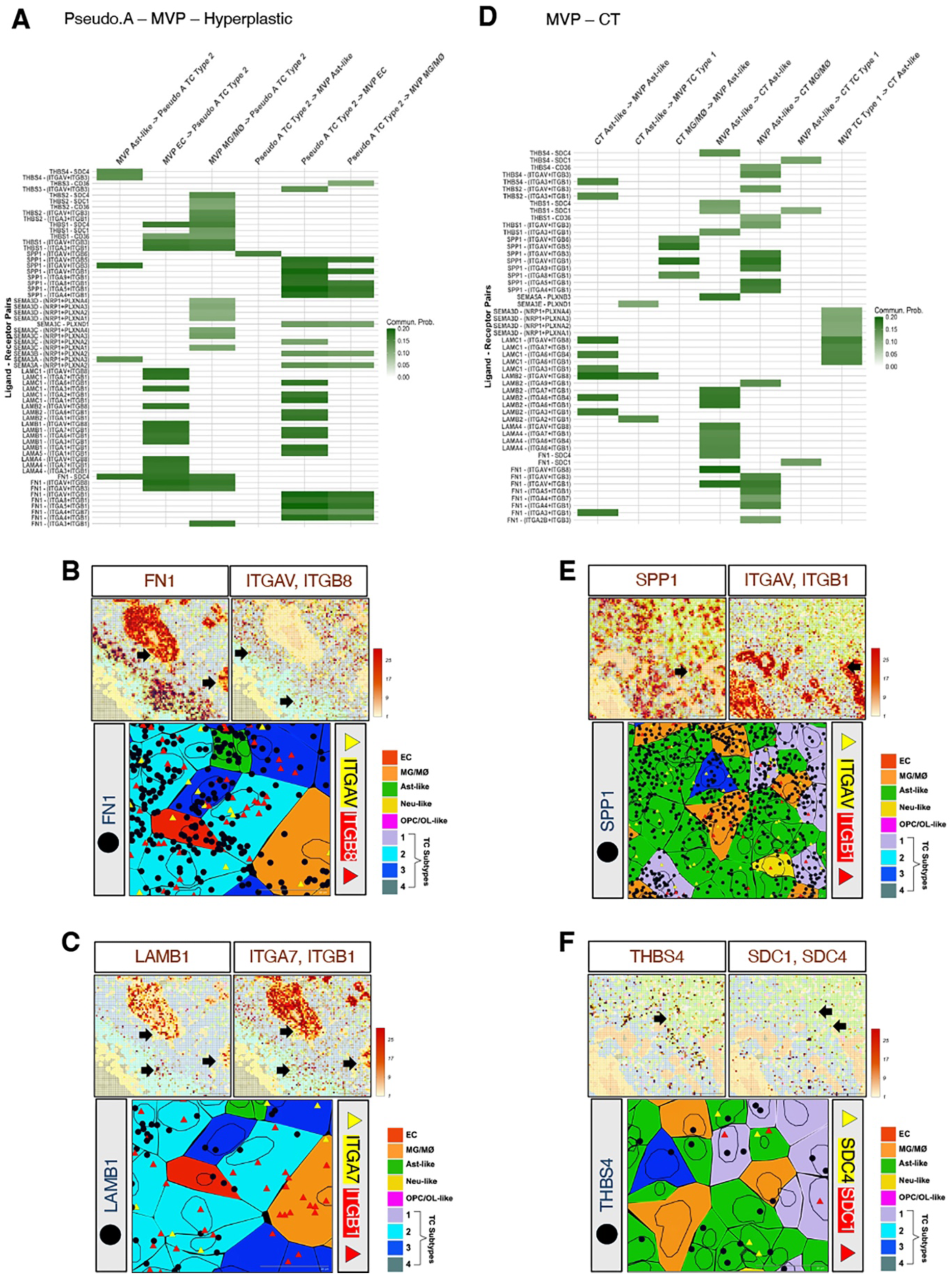
ECM ligand-receptor pairings in different cellular zones in GBM. **(A);** Heatmap showing communication probabilities for top ligand-receptor pairs enriched in TC and stromal cell types localized across the Pseudo A–MVP–Hyperplastic boundary. **(B);** Top panel shows representative transcript heatmaps for FN1 (ligand) and corresponding integrin receptors (ITGAV and ITGB8) across Pseudo.A-MVP-Hyperplastic boundaries. Bottom panel shows high-resolution image displaying spatial localization of transcripts (dark blue circles for FN1 and multi-colored triangles for integrins) in pseudo-colored cell types. Scale bars: 200 µm and 20 µm. **(C);** Top panel shows transcript heatmaps for LAMB1 (ligand) and corresponding integrin receptors (ITGA7 and ITGB1) across Pseudo A–MVP–Hyperplastic boundaries. Bottom panel shows representative high-resolution image displaying spatial localization of the transcripts (dark blue solid circle for LAMB1 and multi-colored triangles for integrins) in UMAP-based cell types (pseudo-colors indicated). Scale bars: 200 µm and 20 µm. **(D);** Heatmap showing communication probabilities for top ligand-receptor pairs enriched in TC and stromal cell types localized across the MVP–CT boundary. **(E);** Top panel shows transcriptomic heatmaps for SPP1 (ligand) and corresponding integrin receptors (ITGAV and ITGB1) across the MVP–CT boundary. Bottom panel shows representative high-resolution image displaying spatial localization of the transcripts (dark blue solid circle for SPP1 and multi-colored triangles for integrins) in UMAP-based cell types (pseudo-colors indicated). Scale bars: 200 µm and 20 µm. **(F);** Top panel shows transcriptomic heatmaps for THBS4 (ligand) and corresponding syndecan receptors (SDC1 and SDC4) across the MVP-CT boundary. Bottom panel shows high-resolution image of transcript spatial localization (dark blue solid circle for THBS4 and multi-colored triangles for syndecans) in pseudocolored cells. Scale bars: 200 µm and 20 µm.

The spatial expression patterns for matrisome and related genes in patient-matched samples from non-cancerous brain and GBM were next analyzed (n=3 of each). The GBM samples contained the tumor leading edge (LE) that included cancer cells infiltrating into brain parenchyma (IT). The annotated LE and IT regions were validated by comparing the expression of gene signatures selected from the Ivy GAP dataset (15). In comparison to the matching non-cancerous brain samples, ECM genes SEMA3E, COL8A1 and IGFBP2 were spatially elevated in the LE/IT regions in the GBM samples (Fig. 7A-B). Some DEGs like FREM2, ANXA2 and GPC4 revealed common enrichment in LE/IT tumor regions across GBM patient samples compared to the overall downregulation of the OPC/OL marker MBP, the neural marker NTRK2, and the ECM glycoprotein EDIL3 (Fig. 7C). While ECM genes SPON1, SDC4 and ANXA1 were elevated in more infiltrating tumor tissue (patients 1 and 2), SEMA3A, SPOCK1 and SDC2 displayed enhanced expression in less infiltrating GBM sample from patient 3 (Fig. 7D-E). The GBM sample from patient 3 exhibited closely positioned LE and IT regions with a distinctively identifiable tumor periphery. Consequently, we performed spatial profiling of these neighboring areas to analyze the differential expression of matrisome genes (Fig. 7F-G). While GFAP, AQP4, GREM1, SOX10 and SEMA3B (selected from top DEGs shown in Fig. 7F) displayed distinct spatial enrichment in the IT region, IGFBP2, NRP1, LGALS1, LGALS3 and CCNB2 (selected from top DEGs shown in Fig. 7F) were elevated in the GBM periphery (Fig. 7G). In comparison to the LE region, more single cells profiled from the IT region revealed higher expression of both GFAP and AQP4 indicating increased reactive astrogliosis in normal brain areas infiltrated by GBM cells (Fig. 7G). Elevated expression for transcripts IGFBP2 and LGALS1 in cells at the tumor periphery (GBM-LE region) indicate aberrant ECM remodeling at the invasive GBM edge (Fig. 7G).

**Figure 7.**
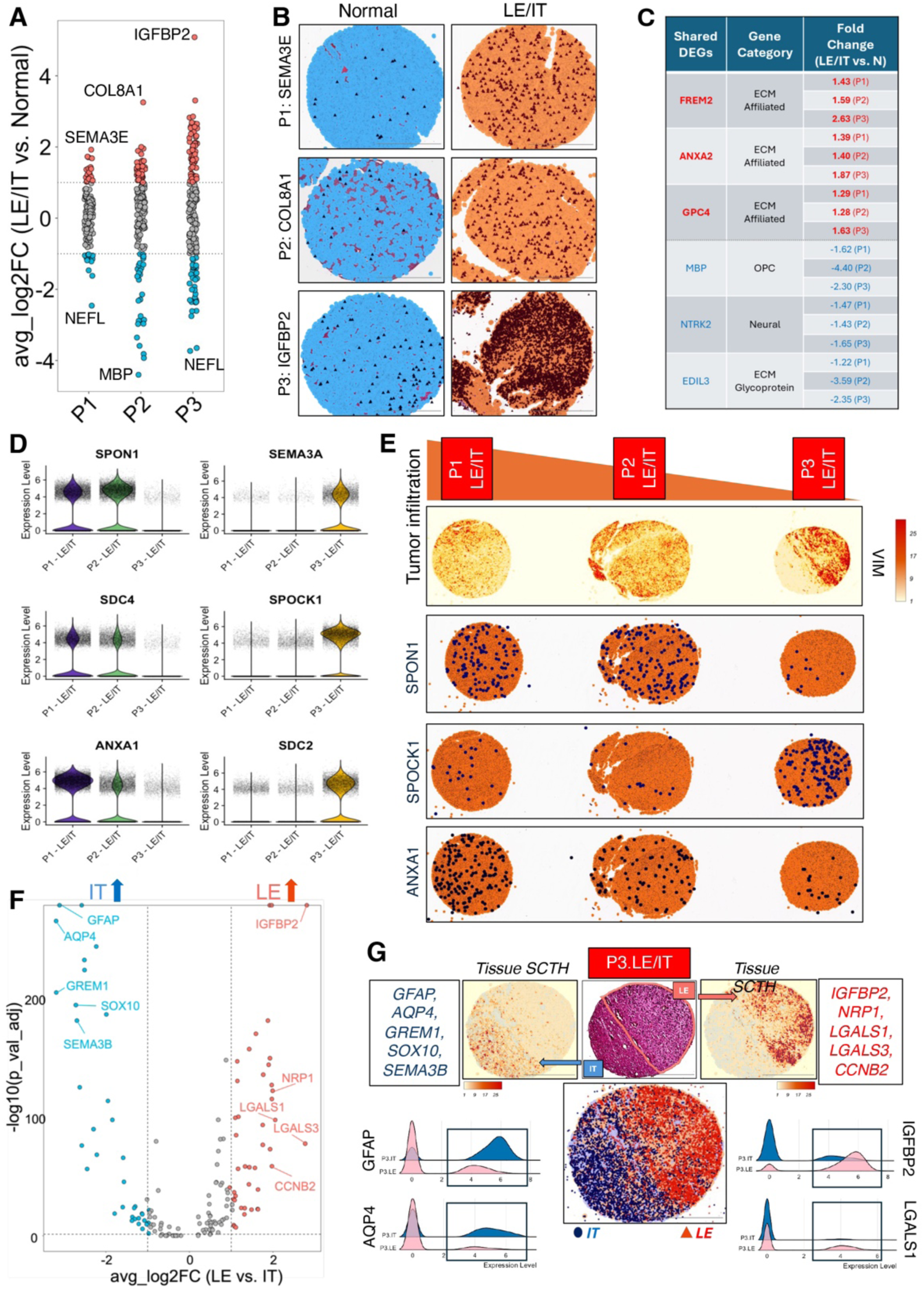
Spatial profiling of matrisome gene expression in GBM infiltrating regions. **(A);** Strip chart showing differentially expressed matrisome genes (upregulated: red and downregulated: blue) in GBM-LE/IT regions compared to matched non-cancerous brain tissue (n=3 patients). **(B);** Representative overall spatial expression of top DEGs upregulated in GBM-LE/IT regions compared to the matched normal samples for individual patients (n=3). Transcripts shown in dark blue (normal) and dark purple (LE/IT) localized in single cells (light blue: normal and light orange: LE/IT). Scale bars: 500 µm. **(C);** Table showing shared upregulated and downregulated DEGs across all three patient GBM-LE/IT samples compared to matched non-cancerous brain tissue. Expression fold-change values in LE/IT samples for shared DEGs for each patient. **(D, E);** Violin plots showing expression of DEGs selected from comparative analysis of GBM-LE/IT patient samples based on degree of infiltration. Representative VIM spatial transcriptomic heatmaps define varying degrees of infiltration across LE/IT samples from three GBM patients (E). Representative images of spatial transcript localization (dark blue) for select genes in single cells (orange) for LE/IT samples across three GBM patients (E). Scale bars: 1000 µm. **(F, G);** Volcano plot showing upregulated DEGs (blue: IT and red: LE) in more distinct GBM-LE and GBM-IT regions for samples from P3 (F). Representative tissue SCTHs showing elevated compartmentalized expression of select DEGs for adjacent IT (left) and LE (right) regions in the same GBM sample from P3 (G). H/E-stained image and spatial image representing cumulative transcript localization of select DEGs for IT (blue) and LE (red) in the GBM sample from P3 are shown in the middle. Ridgeline plots show elevated expression of astroglial markers GFAP and AQP4 in IT region (left) and GBM hallmarks (IGFBP2 and LGALS1) in the LE region t (right). Scale bars: 500 µm.

We profiled matrisome gene expression in a tumor microarray containing punch biopsy samples from grade II diffuse glioma (n=9), grade III astrocytoma (n=10), and grade IV/GBM (n=7) along with non-cancerous brain tissues (n=4) which included some matching samples. All grade II (G.2) and grade III (G.3) samples harbored R132H mutations in the IDH1 gene, whereas the GBM samples (G.4) were wild type for IDH1. Pseudo-bulk sampling identified variance-based distribution of G.2 and G.3 samples when compared to G.4/GBM (Supp. Fig. 8A). Glioma grade-based and integrative single-cell clustering by unsupervised UMAP analyses identified diverse enrichment of TC and multiple stromal cell types in the astrocytoma samples and non-cancerous samples based on differential gene expression profiles (Fig. 8A-D). Specific ECM gene sets were spatially enriched in TC and stromal cell types for each glioma grade (Fig. 8E-J). For example, COL20A1 and SPOCK1 were elevated in TCs for grade II and 3 astrocytoma samples respectively (Fig. 8C-D, F, H), while SPON1 was more diffusely upregulated in grade II tumor and infiltrating multiple stromal cell types. LAMA4 was enriched in tumor ECs in grade III samples (Fig. 8C-D, F, H). For GBM samples, both ANXA2 and POSTN were significantly enhanced in TCs and ECs (Fig. 8C-D, J). The majority of the matrisome genes showing significantly elevated transcript abundance in GBM samples belonged to the ECM glycoprotein and ECM-affiliated gene categories, including ANXA2 (Supp. Fig. 8B). Immunofluorescence labeling detected robust upregulation of annexin-a2 protein expression in GBM versus normal brain as well as lower grade glioma samples (Supp. Fig. 8C).

**Figure 8:**
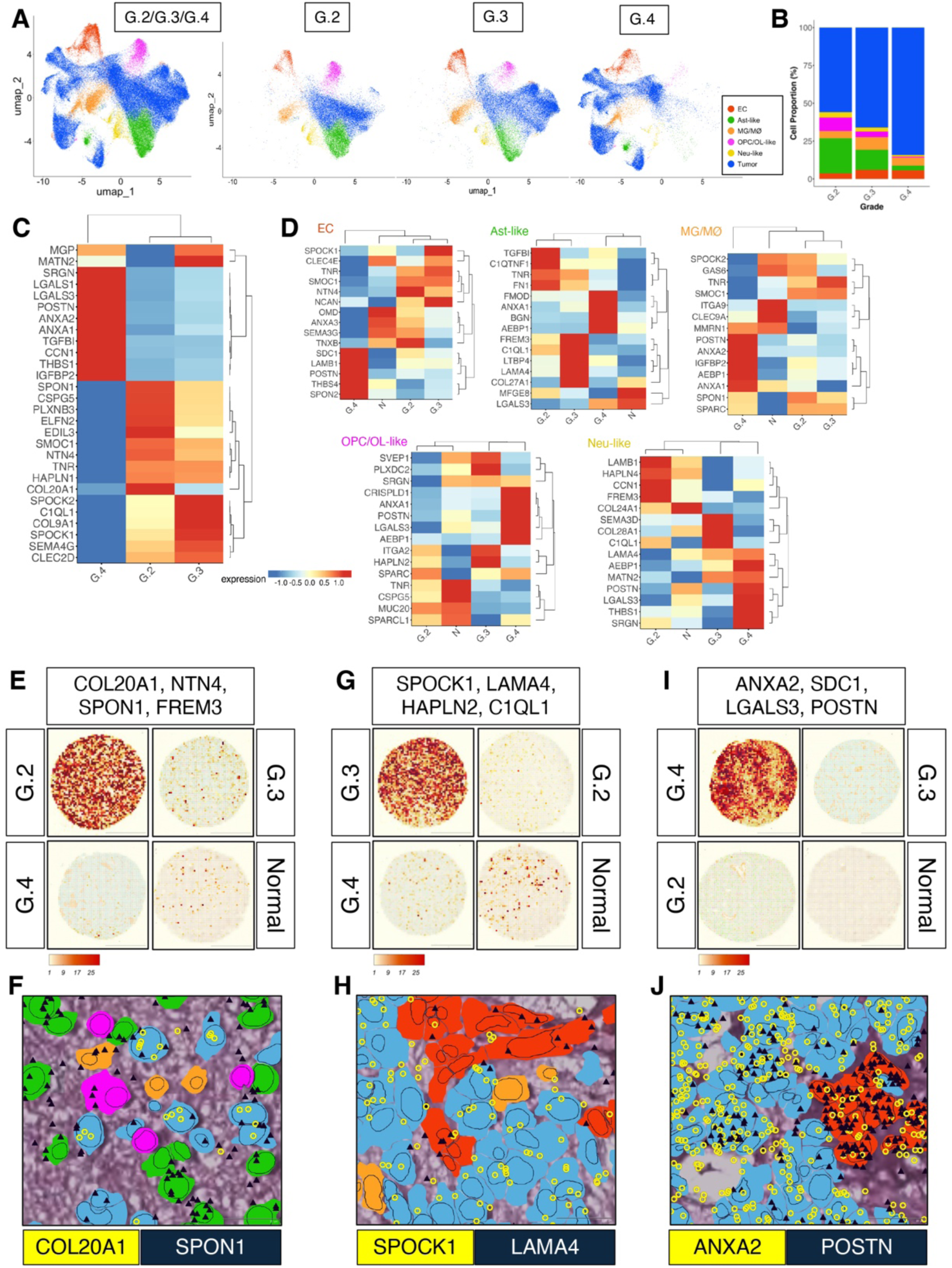
Cross-grade spatial transcriptomic analysis of tumor and stromal cell types. **(A);** UMAP representing unsupervised single-cell combined (left) and grade-separated (right) clustering of tumor and stromal cell types from grade II diffuse glioma (G.2, n=9), grade III astrocytoma (G.3, n=10), and grade IV/GBM (G.4, n=7) samples. Pseudo-colored clusters defined by differential expression profiles for related matrisome markers. **(B);** Stacked bar plot showing comparison of TC and stromal cell-type abundances in tumor samples from different grades (A). Cell-types shown in the plots correspond to assigned pseudo-colors from UMAPs (A). **(C);** Expression heatmap of TCs showing top DEGs across the multiple brain tumor samples from different grades. Expression profiles for tumor cells from G.2 diffuse glioma and G.3 astrocytoma samples show some overlap in comparison to GBM samples that reveal more selective enrichment. Normalized expression values were averaged by tumor grade, log-transformed and scaled. **(D);** Heatmaps of stromal cell-types showing DEGs across the different glioma grades (G.2, G.3 and G.4) in comparison to non-cancerous (N) samples (n=4). Normalized expression values were averaged by sample, log-transformed and scaled. **(E, F);** Representative tissue SCTHs from G.2, G.3, G.4 and non-cancerous samples showing DEGs in G.2 in comparison to G.3, G.4 and normal samples (E). DEGs were selected from single-cell transcriptomic heatmaps for TC and stromal cell-types (C and D) to compare expression levels for matrisome and related gene signatures across brain tumor grades. Representative high-resolution image shows spatial enrichment of COL20A1 transcripts (yellow) in TCs in comparison to higher localization of SPON1 transcripts (dark blue) in surrounding Ast-like cells in G.2 sample. Cells are assigned UMAP-based pseudo-colors shown in A. Scale bars: 500 µm and 20 µm. **(G, H);** SCTHs from G.2, G.3, G.4 and normal samples showing enhanced co-enrichment of DEGs in G.3 in comparison to G.2, G.4 and normal samples (G). DEGs selected from single-cell transcriptomic heatmaps for TC and stromal cell-types shown in C and D to represent and compare expression levels for matrisome and related gene signatures across tumor grades. High-resolution image showing spatial enrichment of SPOCK1 transcripts (yellow) in TCs in comparison to higher localization of LAMA4 transcripts (dark blue) in ECs in the G.3 sample. Cells are assigned UMAP-based pseudo-colors shown in A. Scale bars: 500 µm and 20 µm. **(I, J);** SCTHs from G.2, G.3, G.4 and normal samples showing enhanced co-enrichment of DEGs in G.4 in comparison to G.2, G.3 and normal samples (E). DEGs were selected from single-cell transcriptomic heatmaps for TC and stromal cell-types shown in C and D to represent and compare expression levels for matrisome and related genes. Representative high-resolution image shows concomitant spatial enrichment of both ANXA2 (yellow) and POSTN (dark blue) transcripts in tumor and ECs in G.4 sample. Cells are assigned UMAP-based pseudo-colors shown in A. Scale bars: 500 µm and 20 µm.

## Discussion

Our spatial profiling data showing at least four different tumor cell states largely match with recent single cell transcriptome sequencing studies (17-19). Clusters 2 and 4 in this study localize to hypoxic and necrotic regions, consistent with recent evidence showing hypoxia-dependent ECM remodeling promotes GBM aggressiveness through HIF-1α-dependent mechanisms (20). While matrisome transcripts like ITGA5 and LGALS3 were enriched in TCs and infiltrating stromal cells residing around necrotic regions, the hypoxic tumor regions revealed enhanced expression of IGFBP2 and COL20A1. The ECM-affiliated genes LGALS1 and LGALS3 showed significant upregulation in GBM, particularly in regions of pseudopalisading necrosis. These galectins have recently been implicated in immune evasion and maintenance of cancer stem cells (21), suggesting potential roles in GBM progression and/or therapy resistance.

Spatial enrichment of tumor cluster 1 in the well-vascularized CT region, and cluster 3 in the MVP region suggests involvement in cerebral blood vessel co-option, a process increasingly recognized as crucial for GBM progression (22). In comparison to the CT vasculature, ECM glycoproteins IBSP and MGP, both calcium binding ECM proteins, were upregulated in MVP regions. Studies showing calcium-dependent regulation of GBM invasion (23) suggest that MGP might influence tumor progression through modulation of local calcium dynamics and/or mechanotransduction pathways. In comparison to angiogenic blood vessels in neighboring CT regions, hyperplastic blood vessels within MVP regions differentially expressed matrisome factors like POSTN, OGN and LUM in perivascular MG/MØs. In contrast, SLIT3 and COL8A1 displayed elevated levels in hyperplastic ECs. Interestingly, SLIT3 is a factor that orchestrates angiogenesis during embryonic development (24). Our findings regarding ECM proteoglycans LUM and SRGN build upon emerging evidence that proteoglycans influence the immune microenvironment in GBM (25). The specific enrichment of these factors in ECs and MG/MØs suggests potential roles in the recently described immunosuppressive vascular niche (23). This spatial organization may regulate immune cell trafficking and activation.

Recent studies have demonstrated that integrin signaling influences GBM stem cell maintenance through YAP/TAZ-dependent pathways (26). Our spatial mapping data showing ITGA5 in pseudopalisading TCs and ITGA6/ITGB4 in MVP regions suggests that these signaling events may originate in specific cellular compartments. Our analysis revealed significant upregulation of ANXA1 and ANXA2 in GBM, extending beyond previous TCGA findings by providing crucial spatial contexts (27). While ANXA1 was enriched in MVP, pseudopalisading regions, and more diffusely infiltrative GBM peripheries, ANXA2 was elevated in TCs and ECs in comparison to lower grade tumor samples. The enrichment of these calcium-dependent phospholipid-binding proteins in perivascular regions suggests roles in maintaining the perivascular niche, particularly relevant given recent evidence that annexins regulate calcium-dependent exosome release in GBM (28).

In summary, this work establishes a foundation for several future areas of investigation. First, functional studies are needed to determine how the identified ECM components may cooperatively influence tumor development, progression, and therapy resistance. Second, the integration of our spatial matrisome data with other emerging datasets, including spatial metabolomics (29) and single-cell chromatin accessibility data (30), could provide a more complete understanding of the GBM microenvironment. Finally, the development of therapeutic strategies targeting specific ECM components or their cellular sources will require careful consideration of the spatial organization we have described. Future studies building on this work may lead to more effective treatments for patients with GBM and other brain tumors.

