## Supplementary material for "Single Cell Spatial Profiling Identifies Region-Specific Extracellular Matrix Adhesion and Signaling Networks in Glioblastoma": Supp Data, 8 figures and legends

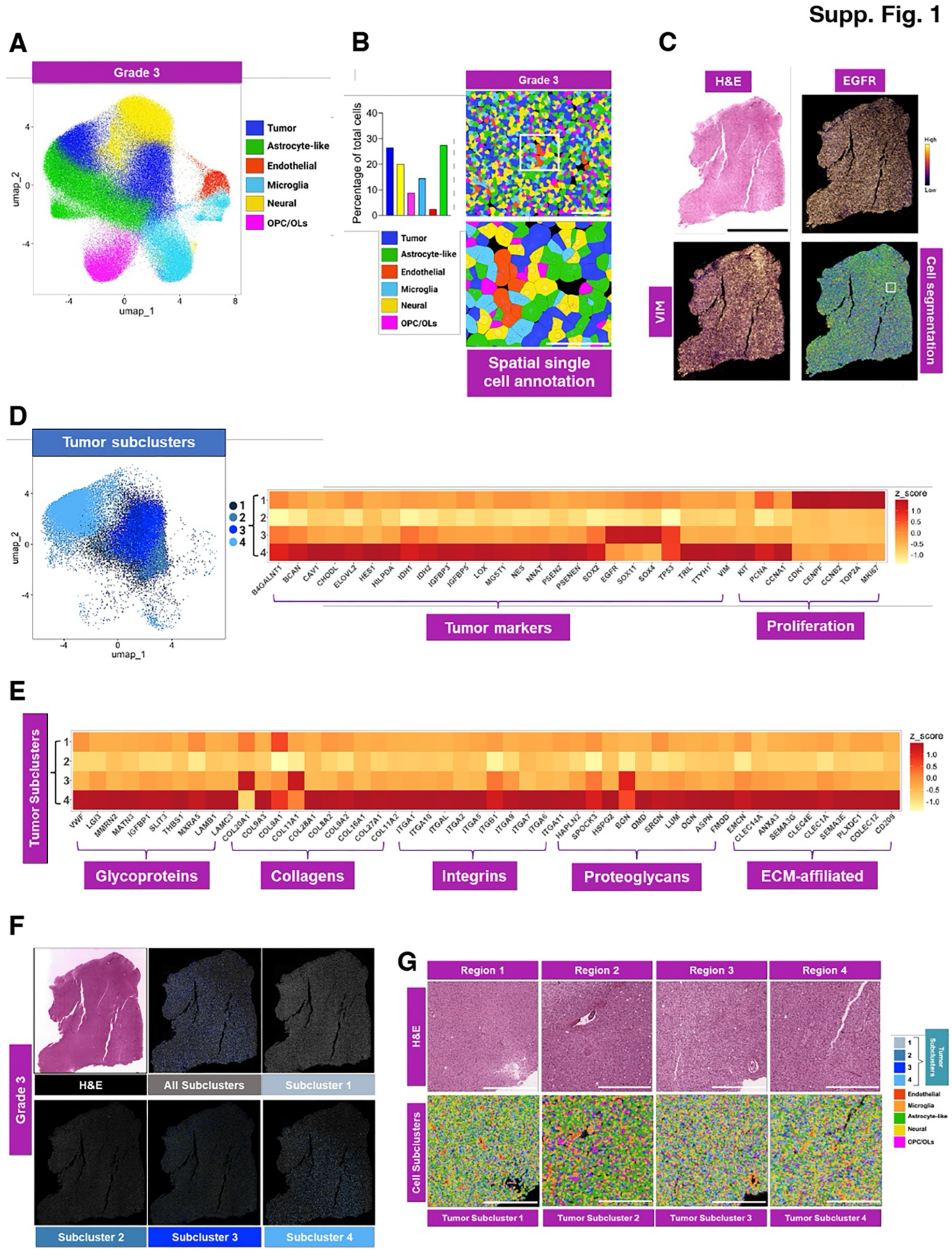

**Supplemental Figure 1. High resolution *in situ* spatial mapping of tumor and matrisome genes in grade III astrocytoma. (A);** UMAP plot of a human grade III astrocytoma sample

(IDH1 R132H) showing pseudo-colored clusters based on matrisome and cell-type specific marker gene expression profiles. **(B);** Graph showing the percentages for the six different stromal and cancer cell types within the grade III sample identified from the UMAP plot (left). Representative image of the grade III astrocytoma sample showing different cell types assigned pseudo-colors (right) based on UMAP clustering (A). Bottom panel shows magnified image of the boxed area with high-resolution cell segmentation and spatial annotation. Scale bars: 200 $\mu$m (upper) and 100 $\mu$m (lower)**. (C);** H&E-stained image of a grade III astrocytoma tissue section (top, left) showing viable tumor. Heatmaps from spatial single cell transcript analysis showing EGFR (top, right), and VIM (bottom, left). Spatial segmentation from UMAP analysis showing annotation of different cell types based on gene enrichment in the viable tumor tissue (bottom, right). Scale bar: 5000 $\mu$m. **(D);** UMAP plot (left) showing four tumor sub-clusters identified by in-depth single cell transcript expression profiling shown in A. Heatmap (right) showing expression profiles of TC and proliferation markers in the four TC sub-clusters. **(E);** Heatmap showing transcript enrichment of select matrisome genes in different TC subtypes. Shown on the horizontal axis are top DEG from each matrisome category. **(F);** Representative H&E-stained section (left panel) and pseudocolored cell images (right panel) for the grade III astrocytoma sample reveals uniform spatial patterning of TC subclusters. Scale bars: 500 $\mu$m. **(G);** Representative H&E (top) and pseudocolored cells (bottom) for four different grade III astrocytoma areas reveals non-distinct and largely overlapping spatial positioning of TCs belonging to the four different subclusters. Scale bars: 500 $\mu$m.

Supp. Fig. 2

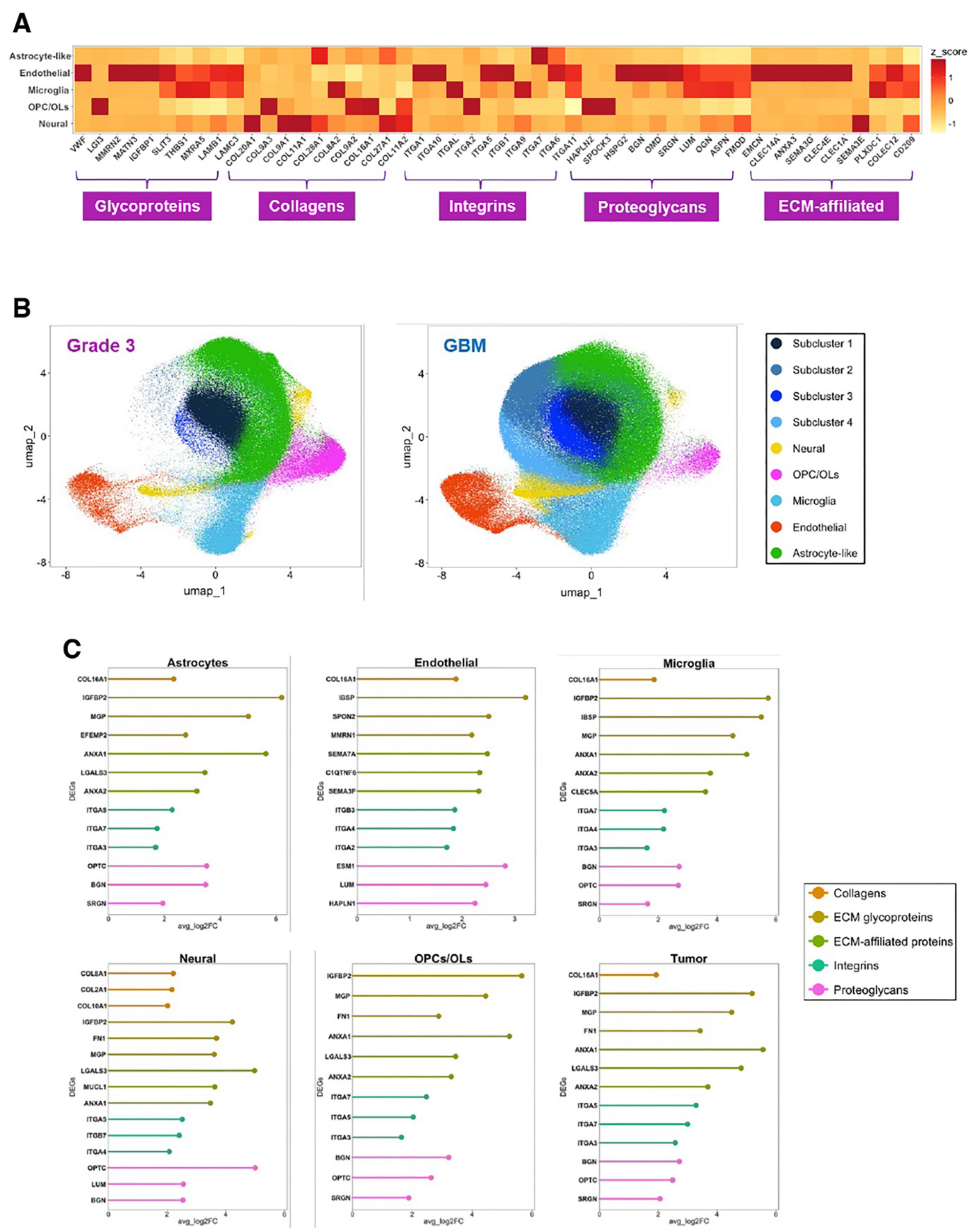

**Supplemental Figure 2. Spatial gene expression analysis of stromal cell types in grade III astrocytoma and GBM. (A);** Heatmap showing enrichment of selected matrisome transcripts in different single cell types in the grade III astrocytoma sample. Shown on the horizontal axis are the top DEGs from each group: ECM glycoproteins, collagens, integrins, proteoglycans, and ECM-affiliated factors. **(B);** Integrated UMAP plots from the human grade III astrocytoma sample (left) and GBM sample (right) showing pseudo-colored cell clusters based on matrisome gene expression profiles. **(C);** Lollipop plots from integrated UMAP subclusters (B) showing transcript expression profiles for different stromal cells and cancer cell clusters in the GBM sample. All DEGs in the lollipop plots are statistically significant ($p < 0.0001$). All transcripts show higher expression in single cells from the GBM versus the grade III astrocytoma.

Supp. Fig. 3

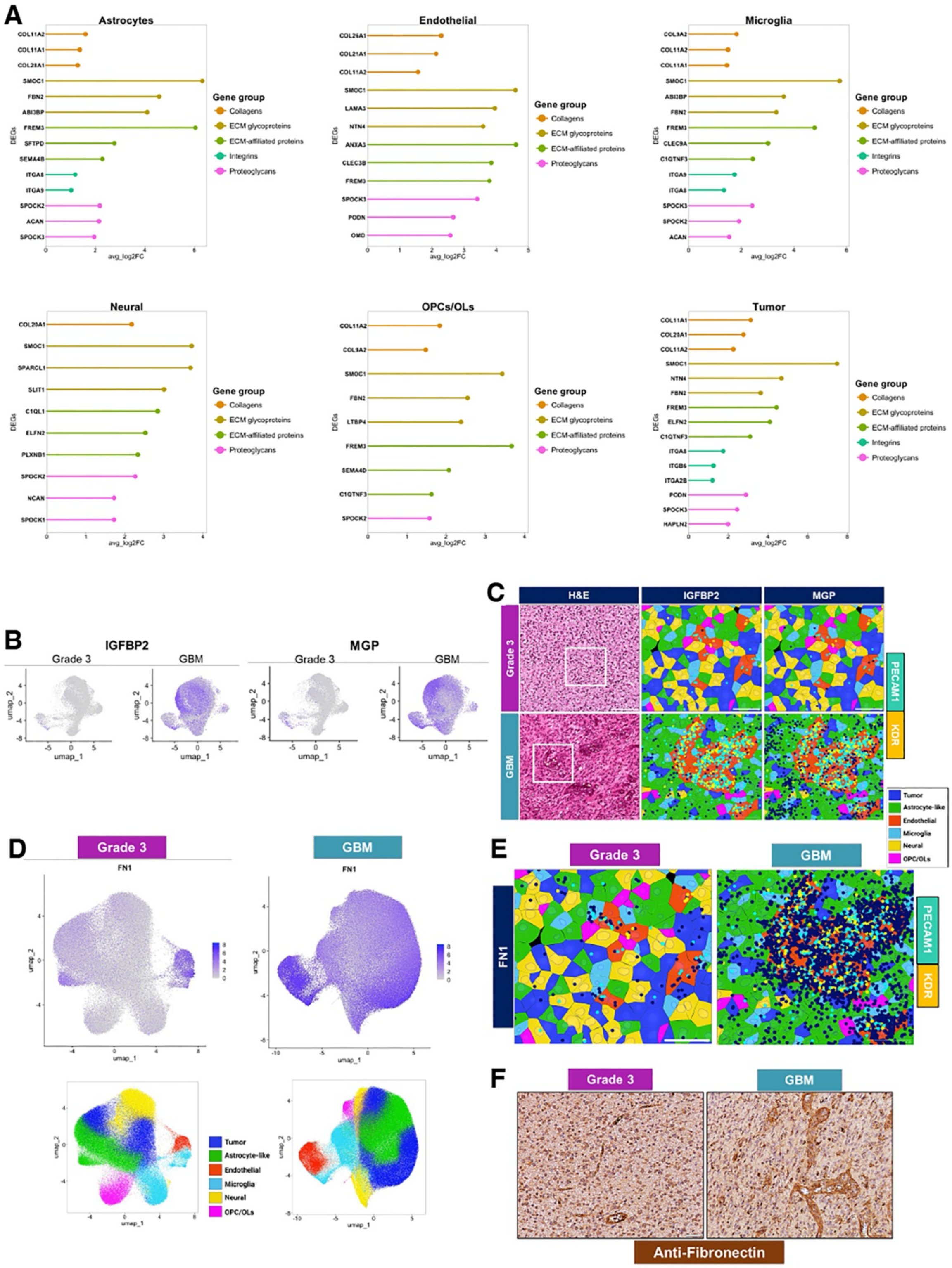

**Supplemental Figure 3. DEGs in grade III astrocytoma sample and spatial mapping of upregulated matrisome genes in GBM. (A);** Single-cell transcript analysis of integrated UMAP cell subclusters, displayed as lollipop plots, showing statistically significant differential gene expression profiles for stromal cells and TC clusters in grade III astrocytoma ($p < 0.0001$). All transcripts display higher expression in cells from the grade III astrocytoma sample versus the GBM sample. **(B);** Integrated UMAP plots showing elevated IGFBP2 (left) and MGP (right) gene expression in multiple cell types in the GBM sample versus the grade III astrocytoma sample. **(C);** Representative images showing *in situ* spatial distribution of the ECM glycoprotein transcripts IGFBP2 and MGP at single-cell resolution. Cells are pseudo-colored based on UMAP analyses of grade III and GBM samples. The right panels are magnified regions from the boxed region of interest (H&E) of the grade III (top) and GBM (bottom) samples. In comparison to grade III astrocytoma, both transcripts show significantly increased expression in different cell types proximal to ECs in the GBM sample. PECAM1; cyan, KDR; yellow and ECM glycoprotein gene transcripts; dark blue. Scale bars: 200 $\mu$m for H&E images and 50 $\mu$m for pseudocolored images. **(D);** UMAP plot for the grade III astrocytoma sample (left) showing FN1 expression primarily in the vascular EC cluster and in some astrocyte-like cells and microglia. This is different than the UMAP plot (right) showing robust FN1 enrichment in multiple cell types in addition to vascular ECs in a GBM samples. The UMAP plots along the bottom show the pseudo-colored cell cluster identities. **(E);** In comparison to grade III astrocytoma (left), there is elevated FN1 mRNA (dark blue transcripts) in GBM (right). **(F);** Representative images showing elevated fibronectin protein expression in GBM by immunohistochemistry. These data validate the spatial transcriptome analysis. Scale bar: 50 $\mu$m.

**Supp. Fig. 4**

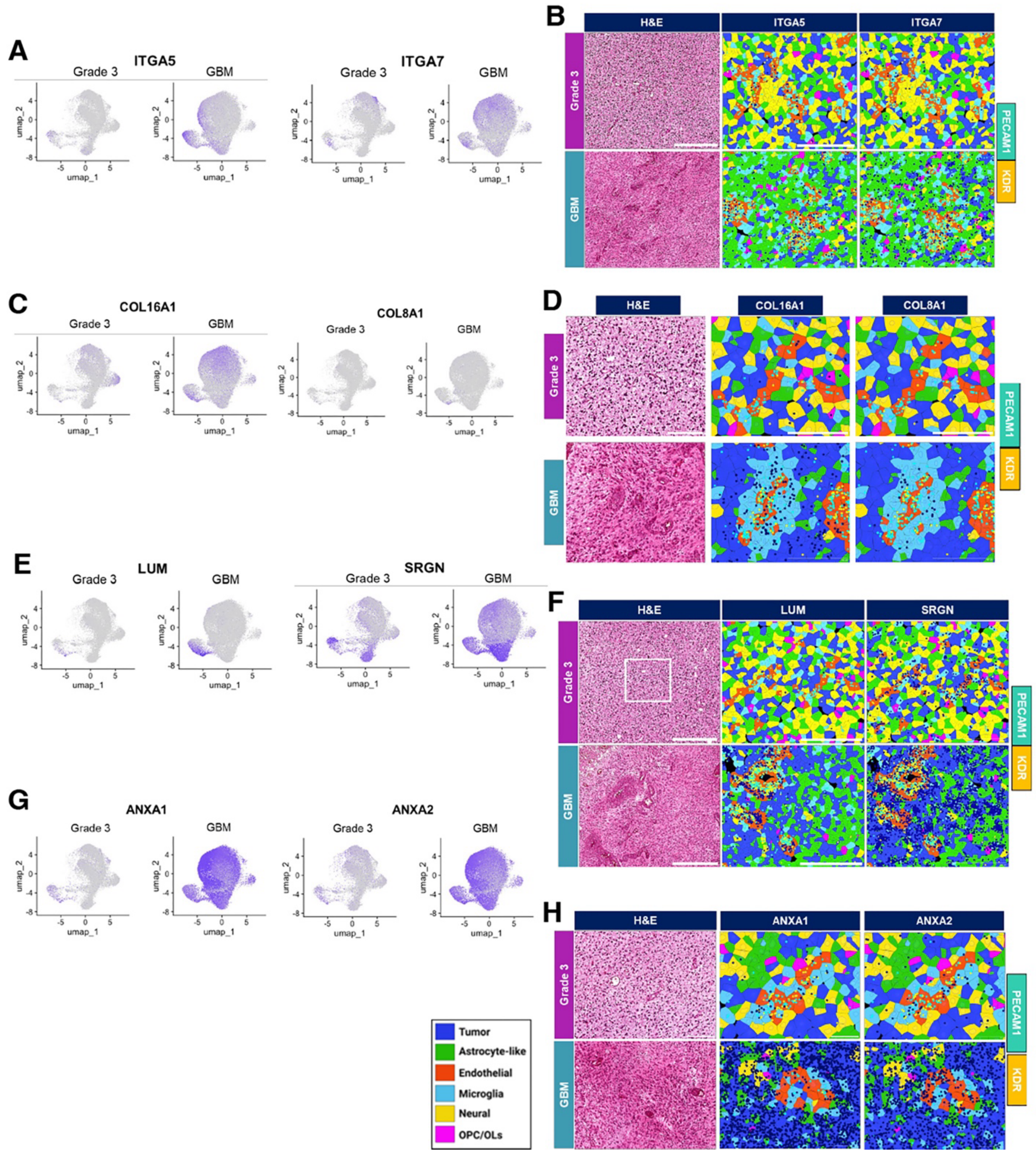

**Supplemental Figure 4. High resolution *in situ* single cell spatial mapping for genes encoding integrins, collagens, proteoglycans and ECM-affiliated factors. (A);** Integrated UMAP plots revealing elevated ITGA5 (left) and ITGA7 (right) transcripts in multiple cell types in the GBM sample versus the grade III astrocytoma sample. **(B);** Representative images showing spatial localization of select integrin transcripts (ITGA5 and ITGA7) at single-cell resolution.

Shown are magnified regions from boxed ROIs (H&E) of the grade III astrocytoma (top) and GBM (bottom). In comparison to grade III sample, ITGA5 transcripts (dark blue) show higher expression in ECs whereas ITGA7 transcripts (dark blue) show elevated expression in perivascular astrocyte-like cells and cancer cells within the GBM sample. PECAM1; cyan, KDR; yellow. Scale bars: 500 $\mu$m for H&E images and 200 $\mu$m for high-resolution single-cell images. **(C);** Integrated UMAP plots revealing elevated COL16A1 (left) and COL8A1 (right) gene expression in multiple cell types in the GBM sample versus the grade III astrocytoma sample. **(D);** Representative images showing spatial distribution of selected collagen transcripts (COL16A1 and COL8A1) at single cell resolution (cells pseudo-colored) in the magnified region from boxed region of interest (H&E) of the grade III astrocytoma (top) and GBM (bottom) tissue sections. In comparison to the grade III tumor, both selected collagen genes show enhanced expression in ECs (red), microglia (light blue), astrocyte-like cells (green), and TCs (dark blue) in the GBM sample. PECAM1; cyan, KDR; yellow and collagen gene transcripts; dark blue. Scale bars: 200 $\mu$m for H&E images and 100 $\mu$m for single cell images. **(E);** Integrated UMAP plots showing elevated LUM (left) and SRGN (right) transcripts in different cell types in the GBM sample versus the grade III astrocytoma sample. Note that LUM expression increases primarily in ECs whereas increased SRGN expression is detected in multiple cell types. **(F);** Representative images showing spatial distribution of proteoglycan transcripts LUM and SRGN at single-cell resolution in the magnified region from boxed ROI (H&E) of the grade III (top) and GBM sample (bottom). In comparison to grade III astrocytoma sample, LUM shows enhanced transcript levels primarily in ECs whereas SRGN shows transcript enrichment in tumor and different stromal cell types in the GBM sample. PECAM1; cyan, KDR; yellow and proteoglycan gene transcripts; dark blue. Scale bars: 500 $\mu$m for H&E images and 200 $\mu$m for pseudocolored images. **(G);** Integrated UMAPs showing elevated ANXA1 (left) and ANXA2 (right) gene expression in different cell types in the GBM sample versus the grade III astrocytoma sample.

**(H);** Representative images showing *in situ* spatial distribution of ECM-affiliated transcripts ANXA1 and ANXA2 at single cell resolution (cells pseudo-colored) in the magnified region from boxed ROI (H&E) of the grade III tissue (top) and GBM (bottom). Both ANXA1 and ANXA2 transcripts show significantly increased expression predominantly in TCs (dark blue) and other cell types: astrocyte-like (green) and microglia (light blue), as well as in ECs (red) in the GBM sample. PECAM1; cyan, KDR; yellow and ECM-affiliated gene transcripts; dark blue. Scale bars: 200 $\mu$m for H&E images and 50 $\mu$m for pseudo-colored images.

Supp. Fig. 5

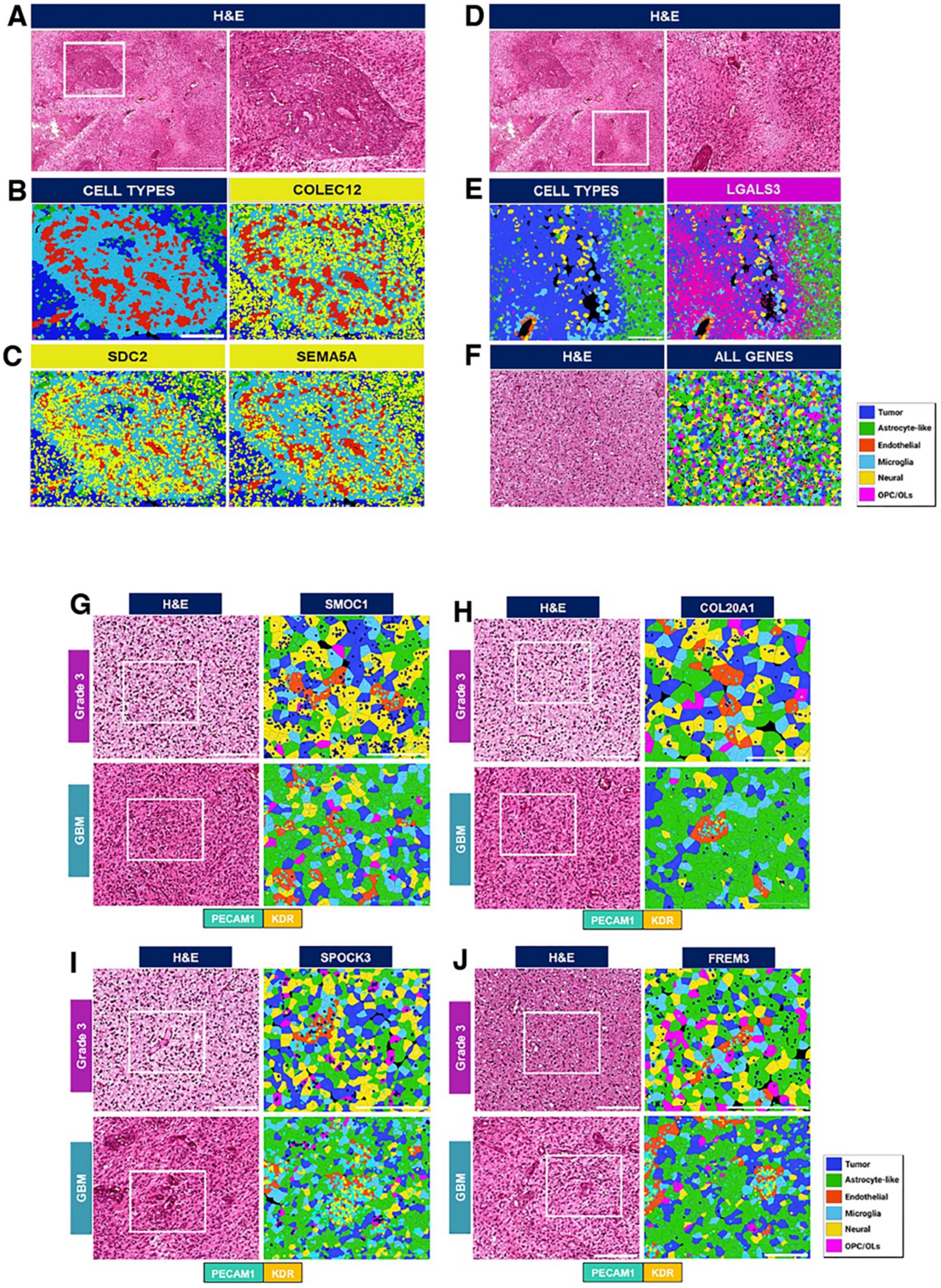

**Supplemental Figure 5. Analysis of matrisome gene expression in regions of neuroinflammation in GBM. (A);** Representative ROI in H&E-stained GBM sample showing perivascular microgliosis. Boxed area on the left is shown as a magnified image on the right.

Scale bars: 100 μm (left) and 200 μm (right). **(B);** Single cell spatial analysis of boxed region shown in (A) identifies clusters of microglial cells (light blue) mixed with ECs (red), (left, B). Note the presence of GBM cells (dark blue) and astrocyte-like cells (green) in the surrounding region. Scale bars: 200 μm. ECM-affiliated transcript COLEC12 (right, B) shows enhanced expression in diverse cell types identified. COLEC12 transcripts are shown in yellow. Scale bars: 200 μm. **(C);** SDC2 (left) and SEMA5A (right) show elevated expression in different cell types localized around necrosis. All gene transcripts are shown in yellow. Scale bars: 200 μm. **(D);** Representative region of interest in H&E-stained GBM sample showing a region with pseudopalisading necrosis. Boxed area on the left is shown as a magnified image on the right. Scale bars: 100 μm (left) and 200 μm (right). **(E);** Single cell transcript analysis of the region shown in (D), highlighting abundant tumor (dark blue) and astrocyte-like (green) cell populations along with less abundant microglia (light blue), neural (yellow) and ECs (red). Increased LGALS3 transcripts (dark pink) in TCs (dark blue) compared to astrocyte-like cells (green) in the pseudopalisading region (right). Scale bar: 200 μm. **(F);** Representative images showing H&E-stained grade III sample (left) and corresponding pseudo-colored cell types (right) display lower levels of transcripts for COLEC12, SDC2, SEMA5A and LGALS3 (all shown in dark blue) in different cell types (right). Scale bars: 200 μm. **(G-J);** Matrisome transcripts with reduced levels in GBM versus grade III astrocytoma. Representative images showing *in situ* single cell spatial transcript localization for grade III (top) and GBM (bottom) from the boxed ROIs shown in corresponding H&E images reveal comparatively elevated cellular transcript enrichment of ECM glycoprotein SMOC1 (G), collagen COL20A1 (H), proteoglycan SPOCK3 (I) and ECM-affiliated gene FREM3 (J) in grade III astrocytoma. *In situ* spatial localization of matrisome transcripts (dark blue) in pseudocolored cells with the EC markers PECAM1 (cyan) and KDR (yellow). Scale bars: 200 μm for H&E images, and 100 μm for high-resolution single-cell spatial transcriptomic images.

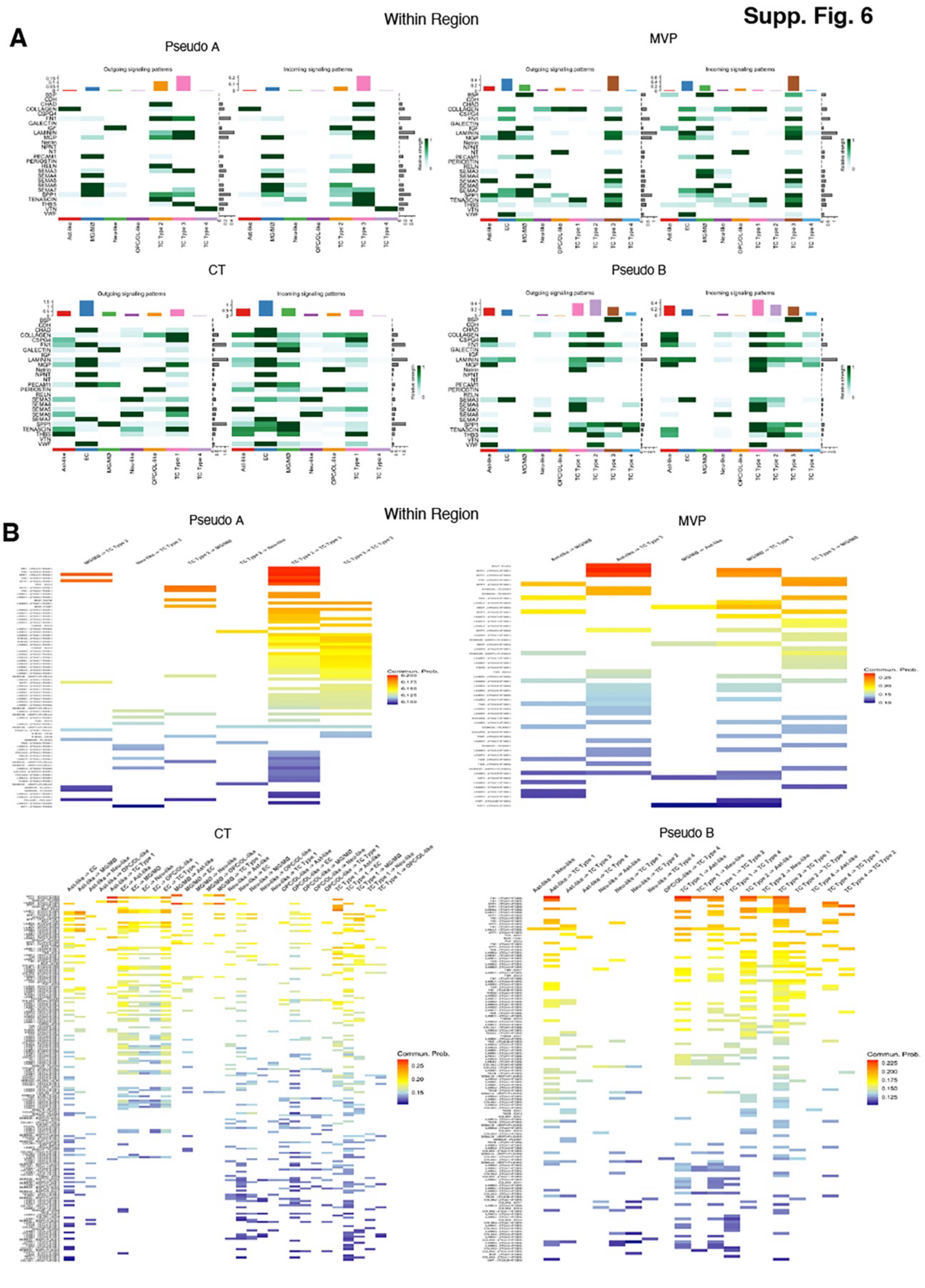

**Supplemental Figure 6. Cell-cell communication networks within GBM anatomic regions.**

**(A);** Heatmap representation of signaling networks showing outgoing and incoming signaling patterns within the Pseudo.A, MVP, CT and Pseudo.B regions. The intensity of each color indicates the strength of ligand-receptor interactions within each pathway, with stronger signals shown in darker green shades. **(B);** Communication probability of all the Ligand-Receptor pairs within regions showing only above-mean communication probability and sorted by their probabilities.

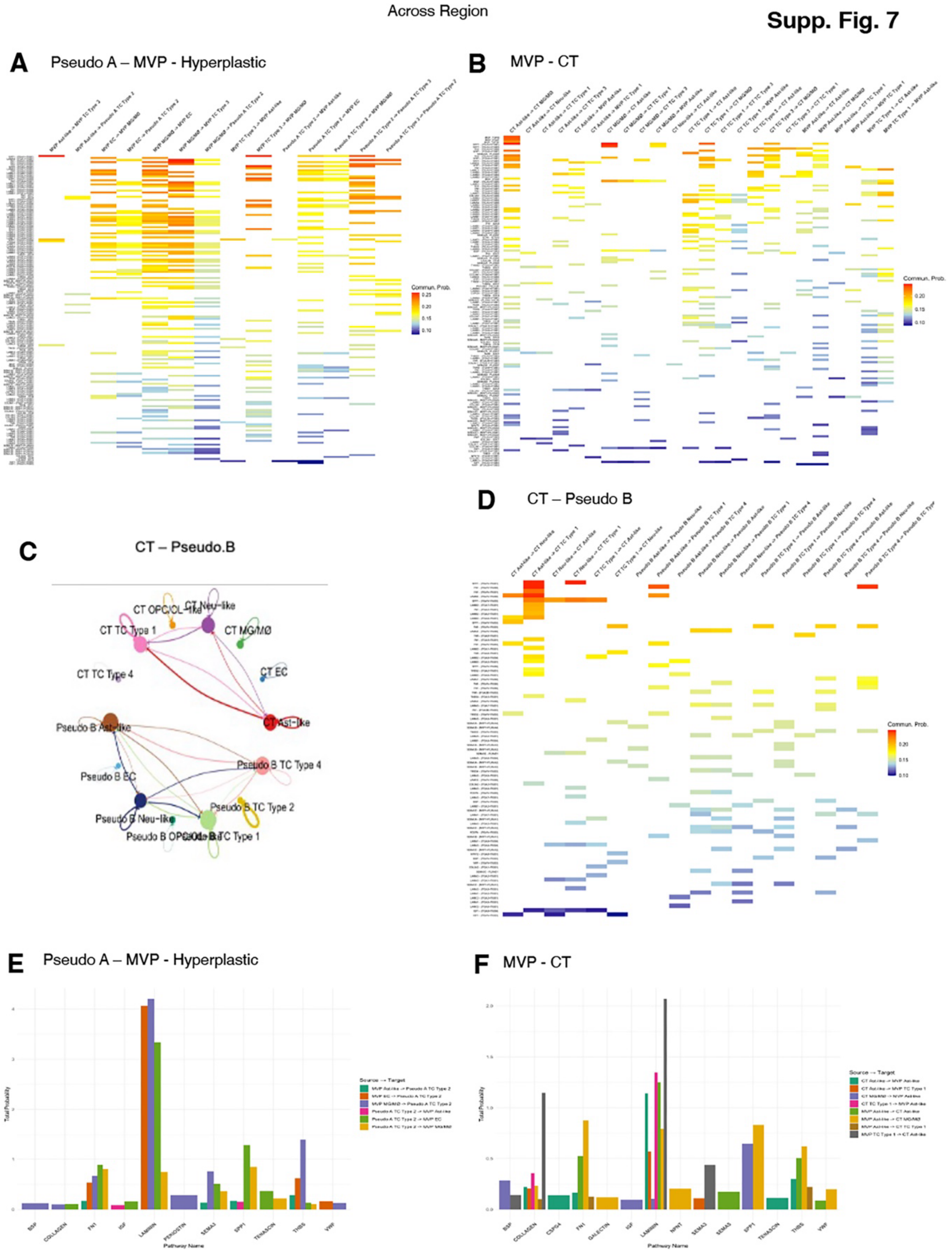

**Supplemental Figure 7. Communication probabilities for signaling pathways inferred by CellChat. (A);** Communication probability of all ligand-receptor pairs across the regions, showing only above-mean communication probability. **(B);** Communication probabilities of all the signaling pathways for Pseudo.A-MVP-Hyperplastic and MVP-CT regions. **(C);** Chord plot summarizing interactions between region-specific cell types across the CT-Pseudo.B boundaries. There was no significant cellular crosstalk detected across the Pseudo.B-CT boundaries. **(D);** Heatmap representation of networks showing outgoing and incoming signaling strength across CT-Pseudo.B boundary. Left panel shows outgoing signals from CT cell types (light green) to recipient Pseudo.B region cell types (dark green). Right panel shows outgoing signals from Pseudo.B region cell types (dark green) to recipient Pseudo.B cell types (light green). **(E);** Bar plot showing distribution of communication probabilities (taken as a sum of all L-R pairs probability) for all the signaling pathways across the region( Pseudo-MVP-Hyperplastic). The bars are colored by cell types showing the directionality of crosstalk across the boundaries. **(F);** Bar plot showing distribution of communication probabilities (taken as a sum of all LR pairs probability) for all the signaling pathways across the region MVP-CT. The bars are colored by cell types showing the directionality of crosstalk across the boundaries.

Supp. Fig. 8

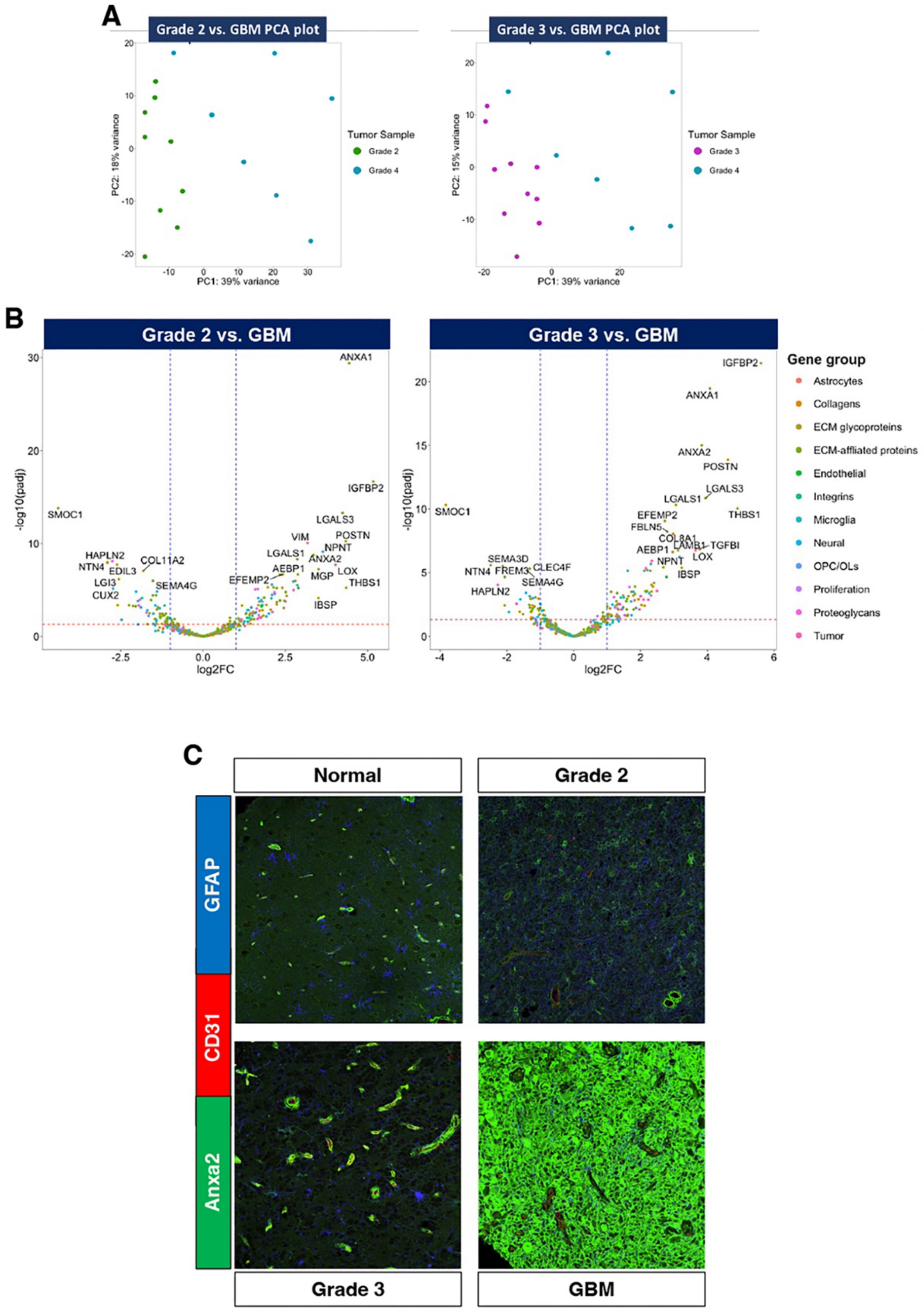

**Supplemental Figure 8. Single cell spatial analysis of matrisome expression in lower grade gliomas and GBM samples/ (A);** PCA plots comparing variance-based distribution of grade II diffuse glioma (n=9) and grade III astrocytoma (n=10) versus GBM (n=7) pseudo-bulk patient samples. **(B);** Volcano plot showing DEGs representing different cell markers and matrisome groups compared across grade II glioma (n=9) versus GBM (n=7) pseudo-bulk samples (left), and grade III astrocytoma (n=10) versus GBM (n=7) pseudo-bulk samples (right) resected from different patients. Selected genes are highlighted that show statistically significant (above horizontal red dotted line) $\log_2$ fold change ≥ +1.0 or ≤ -1.0 (right or left of vertical blue dotted lines respectively) in cellular transcript counts for GBM in comparison to grade II and grade III samples. All analyses were based on pseudo-bulk transcript counts from single cells. **(C);** Expression of annexin-a2 protein was analyzed in fixed sections from non-cancerous brain, grade II diffuse glioma, grade III astrocytoma, and GBM by immunofluorescence. Note that in support of the spatial transcriptomic data for ANXA2 expression (A-B), there is an increase in annexin-a2 protein levels in GBM versus lower grade brain tumor samples.

**Supplemental Table 1. Human brain tumor sample information**. Details of patients with brain tumor grades and IDH mutational status used for in situ single-cell spatial transcriptomic profiling. Information for IDH mutational status is unavailable for one grade 2 glioma sample (*). Another sample from grade 2 glioma is IDH2-mutant (**). The patient/tumor-matched non-cancerous/normal samples are indicated. TMA: tumor microarray, LE/IT: leading edge/infiltrating tumor.

| Tissue Sample Group | Sample ID | Sex | Age (yrs.) | IDH Status | Matched normal samples |
|---|---|---|---|---|---|
| Grade 3 astrocytoma | 806929-6.4-6.18 | F | 33 | Mut | |
| GBM | 865285-6.9-6.23 | F | 54 | WT | |
| TMA 4 GBM (LE/IT) | S-18-028786 | M | 75 | WT | Normal-1 |
| TMA 4 GBM (LE/IT) | S-20-029026 | F | 46 | WT | Normal-2 |
| TMA 4 GBM (LE/IT) | S-20-029629 | F | 71 | WT | Normal-3 |
| TMA 6 Grade 2 | G.2.1 | F | 39 | IDH1 Mut | |
| TMA 6 Grade 2 | G.2.2 | M | 46 | * | |
| TMA 6 Grade 2 | G.2.3 | M | 59 | IDH1 Mut | |
| TMA 6 Grade 2 | G.2.4 | M | 35 | IDH1 Mut | |
| TMA 6 Grade 2 | G.2.6 | M | 27 | IDH1 Mut | |
| TMA 6 Grade 2 | G.2.7 | F | 27 | IDH1 Mut | |
| TMA 6 Grade 2 | G.2.8 | M | 58 | IDH1 Mut | |
| TMA 6 Grade 2 | G.2.9 | F | 26 | IDH2 Mut ** | |
| TMA 6 Grade 2 | G.2.10 | F | 32 | IDH1 Mut | Normal-1 |
| TMA 6 Grade 3 | G.3.1 | F | 27 | IDH1 Mut | |
| TMA 6 Grade 3 | G.3.2 | M | 55 | IDH1 Mut | |
| TMA 6 Grade 3 | G.3.3 | F | 79 | IDH1 Mut | |
| TMA 6 Grade 3 | G.3.4 | F | 53 | IDH1 Mut | |
| TMA 6 Grade 3 | G.3.5 | F | 46 | IDH1 Mut | |
| TMA 6 Grade 3 | G.3.6 | M | 28 | IDH1 Mut | |
| TMA 6 Grade 3 | G.3.7 | M | 55 | IDH1 Mut | |
| TMA 6 Grade 3 | G.3.8 | F | 43 | IDH1 Mut | |
| TMA 6 Grade 3 | G.3.9 | M | 33 | IDH1 Mut | Normal-2 |
| TMA 6 Grade 3 | G.3.10 | M | 52 | IDH1 Mut | Normal-3 |
| TMA 6 Grade 4 | G.4.1 | F | 72 | WT | |
| TMA 6 Grade 4 | G.4.4 | M | 65 | WT | Normal-5 |
| TMA 6 Grade 4 | G.4.5 | F | 57 | WT | |
| TMA 6 Grade 4 | G.4.6 | M | 72 | WT | |
| TMA 6 Grade 4 | G.4.7 | M | 61 | WT | |
| TMA 6 Grade 4 | G.4.8 | M | 47 | WT | |
| TMA 6 Grade 4 | G.4.9 | M | 66 | WT | |

**Supplemental Table 2. Genes in the matrisome probe panel**. Details of the genes (n=478) encoding core ECM factors, ECM-affiliated factors, integrins and other markers enriched in tumor and/or brain stromal cell types, used to design the multi-probe panel to detect the spatial cellular expression and functional aspects of the human matrisome and related molecular signatures. Also included are ensemble IDs and categories (color coded).

| Gene | Ensembl_ID | Category |
|---|---|---|
| ABI3BP | ENSG00000154175 | ECM glycoproteins |
| ADIPOQ | ENSG00000181092 | ECM glycoproteins |
| AEBP1 | ENSG00000106624 | ECM glycoproteins |
| AGRN | ENSG00000188157 | ECM glycoproteins |
| AMELX | ENSG00000125363 | ECM glycoproteins |
| BMPER | ENSG00000164619 | ECM glycoproteins |
| CRELD1 | ENSG00000163703 | ECM glycoproteins |
| CRELD2 | ENSG00000184164 | ECM glycoproteins |
| CRISPLD1 | ENSG00000121005 | ECM glycoproteins |
| CRISPLD2 | ENSG00000103196 | ECM glycoproteins |
| CCN2 | ENSG00000118523 | ECM glycoproteins |
| CYR61 | ENSG00000142871 | ECM glycoproteins |
| DMBT1 | ENSG00000187908 | ECM glycoproteins |
| DPT | ENSG00000143196 | ECM glycoproteins |
| ECM1 | ENSG00000143369 | ECM glycoproteins |
| ECM2 | ENSG00000106823 | ECM glycoproteins |
| EDIL3 | ENSG00000164176 | ECM glycoproteins |
| EFEMP1 | ENSG00000115380 | ECM glycoproteins |
| EFEMP2 | ENSG00000172638 | ECM glycoproteins |
| EGFLAM | ENSG00000164318 | ECM glycoproteins |
| ELN | ENSG00000049540 | ECM glycoproteins |
| FBLN2 | ENSG00000163520 | ECM glycoproteins |
| FBLN5 | ENSG00000140092 | ECM glycoproteins |
| FBLN7 | ENSG00000144152 | ECM glycoproteins |
| FBN1 | ENSG00000166147 | ECM glycoproteins |
| FBN2 | ENSG00000138829 | ECM glycoproteins |
| FBN3 | ENSG00000142449 | ECM glycoproteins |
| FGL1 | ENSG00000104760 | ECM glycoproteins |
| FGL2 | ENSG00000127951 | ECM glycoproteins |
| FN1 | ENSG00000115414 | ECM glycoproteins |
| FNDC1 | ENSG00000164694 | ECM glycoproteins |
| FNDC7 | ENSG00000143107 | ECM glycoproteins |
| FNDC8 | ENSG00000073598 | ECM glycoproteins |
| FRAS1 | ENSG00000138759 | ECM glycoproteins |
| GAS6 | ENSG00000183087 | ECM glycoproteins |

| | | |
|---|---|---|
| IBSP | ENSG00000029559 | ECM glycoproteins |
| IGFALS | ENSG00000099769 | ECM glycoproteins |
| IGFBP1 | ENSG00000146678 | ECM glycoproteins |
| IGFBP2 | ENSG00000115457 | ECM glycoproteins |
| IGFBP6 | ENSG00000167779 | ECM glycoproteins |
| IGFBP7 | ENSG00000163453 | ECM glycoproteins |
| IGFBPL1 | ENSG00000137142 | ECM glycoproteins |
| IGSF10 | ENSG00000152580 | ECM glycoproteins |
| LAMA1 | ENSG00000101680 | ECM glycoproteins |
| LAMA3 | ENSG00000053747 | ECM glycoproteins |
| LAMA4 | ENSG00000112769 | ECM glycoproteins |
| LAMA5 | ENSG00000130702 | ECM glycoproteins |
| LAMB1 | ENSG00000091136 | ECM glycoproteins |
| LAMB2 | ENSG00000172037 | ECM glycoproteins |
| LAMB3 | ENSG00000196878 | ECM glycoproteins |
| LAMB4 | ENSG00000091128 | ECM glycoproteins |
| LAMC1 | ENSG00000135862 | ECM glycoproteins |
| LAMC2 | ENSG00000058085 | ECM glycoproteins |
| LAMC3 | ENSG00000050555 | ECM glycoproteins |
| LGI1 | ENSG00000108231 | ECM glycoproteins |
| LGI2 | ENSG00000153012 | ECM glycoproteins |
| LGI3 | ENSG00000168481 | ECM glycoproteins |
| LGI4 | ENSG00000153902 | ECM glycoproteins |
| LRG1 | ENSG00000171236 | ECM glycoproteins |
| LTBP1 | ENSG00000049323 | ECM glycoproteins |
| LTBP2 | ENSG00000119681 | ECM glycoproteins |
| LTBP3 | ENSG00000168056 | ECM glycoproteins |
| LTBP4 | ENSG00000090006 | ECM glycoproteins |
| MATN2 | ENSG00000132561 | ECM glycoproteins |
| MATN3 | ENSG00000132031 | ECM glycoproteins |
| MFGE8 | ENSG00000140545 | ECM glycoproteins |
| MGP | ENSG00000111341 | ECM glycoproteins |
| MMRN1 | ENSG00000138722 | ECM glycoproteins |
| MMRN2 | ENSG00000173269 | ECM glycoproteins |
| MXRA5 | ENSG00000101825 | ECM glycoproteins |
| NDNF | ENSG00000173376 | ECM glycoproteins |
| NID1 | ENSG00000116962 | ECM glycoproteins |
| NOV | ENSG00000136999 | ECM glycoproteins |
| NTN1 | ENSG00000065320 | ECM glycoproteins |
| NTN3 | ENSG00000162068 | ECM glycoproteins |
| NTN4 | ENSG00000074527 | ECM glycoproteins |
| NTN5 | ENSG00000142233 | ECM glycoproteins |
| PAPLN | ENSG00000100767 | ECM glycoproteins |

| | | |
|---|---|---|
| POSTN | ENSG00000133110 | ECM glycoproteins |
| RELN | ENSG00000189056 | ECM glycoproteins |
| RSPO1 | ENSG00000169218 | ECM glycoproteins |
| RSPO2 | ENSG00000147655 | ECM glycoproteins |
| RSPO3 | ENSG00000146374 | ECM glycoproteins |
| RSPO4 | ENSG00000101282 | ECM glycoproteins |
| SLIT1 | ENSG00000187122 | ECM glycoproteins |
| SLIT2 | ENSG00000145147 | ECM glycoproteins |
| SLIT3 | ENSG00000184347 | ECM glycoproteins |
| SMOC1 | ENSG00000198732 | ECM glycoproteins |
| SNED1 | ENSG00000162804 | ECM glycoproteins |
| SPARC | ENSG00000113140 | ECM glycoproteins |
| SPARCL1 | ENSG00000152583 | ECM glycoproteins |
| SPON1 | ENSG00000262655 | ECM glycoproteins |
| SPON2 | ENSG00000159674 | ECM glycoproteins |
| SPP1 | ENSG00000118785 | ECM glycoproteins |
| SVEP1 | ENSG00000165124 | ECM glycoproteins |
| TGFBI | ENSG00000120708 | ECM glycoproteins |
| THBS1 | ENSG00000137801 | ECM glycoproteins |
| THBS2 | ENSG00000186340 | ECM glycoproteins |
| THBS3 | ENSG00000169231 | ECM glycoproteins |
| THBS4 | ENSG00000113296 | ECM glycoproteins |
| TNR | ENSG00000116147 | ECM glycoproteins |
| TNXB | ENSG00000168477 | ECM glycoproteins |
| VTN | ENSG00000109072 | ECM glycoproteins |
| VWF | ENSG00000110799 | ECM glycoproteins |
| COL10A1 | ENSG00000123500 | Collagens |
| COL11A1 | ENSG00000060718 | Collagens |
| COL11A2 | ENSG00000204248 | Collagens |
| COL16A1 | ENSG00000084636 | Collagens |
| COL17A1 | ENSG00000065618 | Collagens |
| COL20A1 | ENSG00000101203 | Collagens |
| COL21A1 | ENSG00000124749 | Collagens |
| COL23A1 | ENSG00000050767 | Collagens |
| COL24A1 | ENSG00000171502 | Collagens |
| COL26A1 | ENSG00000160963 | Collagens |
| COL27A1 | ENSG00000196739 | Collagens |
| COL28A1 | ENSG00000215018 | Collagens |
| COL2A1 | ENSG00000139219 | Collagens |
| COL7A1 | ENSG00000114270 | Collagens |
| COL8A1 | ENSG00000144810 | Collagens |
| COL8A2 | ENSG00000171812 | Collagens |

| | | |
|---|---|---|
| COL9A1 | ENSG00000112280 | Collagens |
| COL9A2 | ENSG00000049089 | Collagens |
| COL9A3 | ENSG00000092758 | Collagens |
| ACAN | ENSG00000157766 | Proteoglycans |
| ASPN | ENSG00000106819 | Proteoglycans |
| BGN | ENSG00000182492 | Proteoglycans |
| CHAD | ENSG00000136457 | Proteoglycans |
| ESM1 | ENSG00000164283 | Proteoglycans |
| FMOD | ENSG00000122176 | Proteoglycans |
| HAPLN1 | ENSG00000145681 | Proteoglycans |
| HAPLN2 | ENSG00000132702 | Proteoglycans |
| HAPLN3 | ENSG00000140511 | Proteoglycans |
| HAPLN4 | ENSG00000187664 | Proteoglycans |
| HSPG2 | ENSG00000142798 | Proteoglycans |
| IMPG1 | ENSG00000112706 | Proteoglycans |
| IMPG2 | ENSG00000081148 | Proteoglycans |
| LUM | ENSG00000139329 | Proteoglycans |
| NCAN | ENSG00000130287 | Proteoglycans |
| OGN | ENSG00000106809 | Proteoglycans |
| OMD | ENSG00000127083 | Proteoglycans |
| OPTC | ENSG00000188770 | Proteoglycans |
| PODN | ENSG00000174348 | Proteoglycans |
| PODNL1 | ENSG00000132000 | Proteoglycans |
| PRG4 | ENSG00000116690 | Proteoglycans |
| SPOCK1 | ENSG00000152377 | Proteoglycans |
| SPOCK2 | ENSG00000107742 | Proteoglycans |
| SPOCK3 | ENSG00000196104 | Proteoglycans |
| SRGN | ENSG00000122862 | Proteoglycans |
| FREM1 | ENSG00000164946 | ECM-affliated proteins |
| FREM2 | ENSG00000150893 | ECM-affliated proteins |
| FREM3 | ENSG00000183090 | ECM-affliated proteins |
| ELFN1 | ENSG00000225968 | ECM-affliated proteins |
| ELFN2 | ENSG00000166897 | ECM-affliated proteins |
| ELFN2 | ENSG00000243902 | ECM-affliated proteins |
| EMCN | ENSG00000164035 | ECM-affliated proteins |
| FCN1 | ENSG00000085265 | ECM-affliated proteins |
| FCN2 | ENSG00000160339 | ECM-affliated proteins |
| FCN3 | ENSG00000142748 | ECM-affliated proteins |
| SDC1 | ENSG00000115884 | ECM-affliated proteins |
| SDC2 | ENSG00000169439 | ECM-affliated proteins |
| SDC3 | ENSG00000162512 | ECM-affliated proteins |

| | | |
|---|---|---|
| SDC4 | ENSG00000124145 | ECM-affliated proteins |
| SEMA3A | ENSG00000075213 | ECM-affliated proteins |
| SEMA3B | ENSG00000012171 | ECM-affliated proteins |
| SEMA3C | ENSG00000075223 | ECM-affliated proteins |
| SEMA3D | ENSG00000153993 | ECM-affliated proteins |
| SEMA3E | ENSG00000170381 | ECM-affliated proteins |
| SEMA3F | ENSG00000001617 | ECM-affliated proteins |
| SEMA3G | ENSG00000010319 | ECM-affliated proteins |
| SEMA4A | ENSG00000196189 | ECM-affliated proteins |
| SEMA4B | ENSG00000185033 | ECM-affliated proteins |
| SEMA4C | ENSG00000168758 | ECM-affliated proteins |
| SEMA4D | ENSG00000187764 | ECM-affliated proteins |
| SEMA4F | ENSG00000135622 | ECM-affliated proteins |
| SEMA4G | ENSG00000095539 | ECM-affliated proteins |
| SEMA5A | ENSG00000112902 | ECM-affliated proteins |
| SEMA5B | ENSG00000082684 | ECM-affliated proteins |
| SEMA6A | ENSG00000092421 | ECM-affliated proteins |
| SEMA6B | ENSG00000167680 | ECM-affliated proteins |
| SEMA6C | ENSG00000143434 | ECM-affliated proteins |
| SEMA6D | ENSG00000137872 | ECM-affliated proteins |
| SEMA7A | ENSG00000138623 | ECM-affliated proteins |
| PLXDC1 | ENSG00000161381 | ECM-affliated proteins |
| PLXDC2 | ENSG00000120594 | ECM-affliated proteins |
| PLXNA1 | ENSG00000114554 | ECM-affliated proteins |
| PLXNA2 | ENSG00000076356 | ECM-affliated proteins |
| PLXNA3 | ENSG00000130827 | ECM-affliated proteins |
| PLXNA4 | ENSG00000221866 | ECM-affliated proteins |
| PLXNB1 | ENSG00000164050 | ECM-affliated proteins |
| PLXNB2 | ENSG00000196576 | ECM-affliated proteins |
| PLXNB3 | ENSG00000198753 | ECM-affliated proteins |
| PLXNC1 | ENSG00000136040 | ECM-affliated proteins |
| PLXND1 | ENSG00000004399 | ECM-affliated proteins |
| MUC1 | ENSG00000185499 | ECM-affliated proteins |
| MUC12 | ENSG00000205277 | ECM-affliated proteins |
| MUC13 | ENSG00000173702 | ECM-affliated proteins |
| MUC15 | ENSG00000169550 | ECM-affliated proteins |
| MUC16 | ENSG00000181143 | ECM-affliated proteins |
| MUC17 | ENSG00000169876 | ECM-affliated proteins |
| MUC19 | ENSG00000205592 | ECM-affliated proteins |
| MUC2 | ENSG00000198788 | ECM-affliated proteins |
| MUC20 | ENSG00000176945 | ECM-affliated proteins |
| MUC21 | ENSG00000204544 | ECM-affliated proteins |
| MUC22 | ENSG00000261272 | ECM-affliated proteins |

| | | |
|---|---|---|
| MUC3A | ENSG00000169894 | ECM-affliated proteins |
| MUC4 | ENSG00000145113 | ECM-affliated proteins |
| MUC5AC | ENSG00000215182 | ECM-affliated proteins |
| MUC5B | ENSG00000117983 | ECM-affliated proteins |
| MUC6 | ENSG00000184956 | ECM-affliated proteins |
| MUC7 | ENSG00000171195 | ECM-affliated proteins |
| MUCL1 | ENSG00000172551 | ECM-affliated proteins |
| GPC1 | ENSG00000063660 | ECM-affliated proteins |
| GPC2 | ENSG00000213420 | ECM-affliated proteins |
| GPC3 | ENSG00000147257 | ECM-affliated proteins |
| GPC4 | ENSG00000076716 | ECM-affliated proteins |
| GPC5 | ENSG00000179399 | ECM-affliated proteins |
| GPC6 | ENSG00000183098 | ECM-affliated proteins |
| SFTA2 | ENSG00000196260 | ECM-affliated proteins |
| SFTA3 | ENSG00000229415 | ECM-affliated proteins |
| SFTPA1 | ENSG00000122852 | ECM-affliated proteins |
| SFTPA2 | ENSG00000185303 | ECM-affliated proteins |
| SFTPB | ENSG00000168878 | ECM-affliated proteins |
| SFTPC | ENSG00000168484 | ECM-affliated proteins |
| SFTPD | ENSG00000133661 | ECM-affliated proteins |
| CD209 | ENSG00000090659 | ECM-affliated proteins |
| CLC | ENSG00000105205 | ECM-affliated proteins |
| CLEC10A | ENSG00000132514 | ECM-affliated proteins |
| CLEC12A | ENSG00000172322 | ECM-affliated proteins |
| CLEC12B | ENSG00000256660 | ECM-affliated proteins |
| CLEC14A | ENSG00000176435 | ECM-affliated proteins |
| CLEC17A | ENSG00000187912 | ECM-affliated proteins |
| CLEC18A | ENSG00000157322 | ECM-affliated proteins |
| CLEC18B | ENSG00000140839 | ECM-affliated proteins |
| CLEC18C | ENSG00000157335 | ECM-affliated proteins |
| CLEC19A | ENSG00000261210 | ECM-affliated proteins |
| CLEC1A | ENSG00000150048 | ECM-affliated proteins |
| CLEC1B | ENSG00000165682 | ECM-affliated proteins |
| CLEC2A | ENSG00000188393 | ECM-affliated proteins |
| CLEC2B | ENSG00000110852 | ECM-affliated proteins |
| CLEC2D | ENSG00000069493 | ECM-affliated proteins |
| CLEC2L | ENSG00000236279 | ECM-affliated proteins |
| CLEC3A | ENSG00000166509 | ECM-affliated proteins |
| CLEC3B | ENSG00000163815 | ECM-affliated proteins |
| CLEC4A | ENSG00000111729 | ECM-affliated proteins |
| CLEC4C | ENSG00000198178 | ECM-affliated proteins |
| CLEC4D | ENSG00000166527 | ECM-affliated proteins |
| CLEC4E | ENSG00000166523 | ECM-affliated proteins |

| | | |
|---|---|---|
| CLEC4F | ENSG00000152672 | ECM-affliated proteins |
| CLEC4G | ENSG00000182566 | ECM-affliated proteins |
| CLEC4M | ENSG00000104938 | ECM-affliated proteins |
| CLEC5A | ENSG00000258227 | ECM-affliated proteins |
| CLEC6A | ENSG00000205846 | ECM-affliated proteins |
| CLEC7A | ENSG00000172243 | ECM-affliated proteins |
| CLEC9A | ENSG00000197992 | ECM-affliated proteins |
| COLEC10 | ENSG00000184374 | ECM-affliated proteins |
| COLEC11 | ENSG00000118004 | ECM-affliated proteins |
| COLEC12 | ENSG00000158270 | ECM-affliated proteins |
| CSPG4 | ENSG00000173546 | ECM-affliated proteins |
| CSPG5 | ENSG00000114646 | ECM-affliated proteins |
| C1QA | ENSG00000173372 | ECM-affliated proteins |
| C1QB | ENSG00000173369 | ECM-affliated proteins |
| C1QC | ENSG00000159189 | ECM-affliated proteins |
| C1QL1 | ENSG00000131094 | ECM-affliated proteins |
| C1QL2 | ENSG00000144119 | ECM-affliated proteins |
| C1QL3 | ENSG00000165985 | ECM-affliated proteins |
| C1QL4 | ENSG00000186897 | ECM-affliated proteins |
| C1QTNF1 | ENSG00000173918 | ECM-affliated proteins |
| C1QTNF2 | ENSG00000145861 | ECM-affliated proteins |
| C1QTNF3 | ENSG00000082196 | ECM-affliated proteins |
| C1QTNF5 | ENSG00000223953 | ECM-affliated proteins |
| C1QTNF6 | ENSG00000133466 | ECM-affliated proteins |
| C1QTNF7 | ENSG00000163145 | ECM-affliated proteins |
| C1QTNF8 | ENSG00000184471 | ECM-affliated proteins |
| C1QTNF9 | ENSG00000240654 | ECM-affliated proteins |
| ANXA1 | ENSG00000135046 | ECM-affliated proteins |
| ANXA10 | ENSG00000109511 | ECM-affliated proteins |
| ANXA11 | ENSG00000122359 | ECM-affliated proteins |
| ANXA13 | ENSG00000104537 | ECM-affliated proteins |
| ANXA2 | ENSG00000182718 | ECM-affliated proteins |
| ANXA3 | ENSG00000138772 | ECM-affliated proteins |
| ANXA4 | ENSG00000196975 | ECM-affliated proteins |
| ANXA5 | ENSG00000164111 | ECM-affliated proteins |
| ANXA6 | ENSG00000197043 | ECM-affliated proteins |
| ANXA7 | ENSG00000138279 | ECM-affliated proteins |
| ANXA8 | ENSG00000265190 | ECM-affliated proteins |
| ANXA8L1 | ENSG00000264230 | ECM-affliated proteins |
| ANXA9 | ENSG00000143412 | ECM-affliated proteins |
| GREM1 | ENSG00000166923 | ECM-affliated proteins |
| HPX | ENSG00000110169 | ECM-affliated proteins |
| LGALSL | ENSG00000119862 | ECM-affliated proteins |

| | | |
|---|---|---|
| ITLN1 | ENSG00000179914 | ECM-affliated proteins |
| ITLN2 | ENSG00000158764 | ECM-affliated proteins |
| LGALS1 | ENSG00000100097 | ECM-affliated proteins |
| LGALS12 | ENSG00000133317 | ECM-affliated proteins |
| LGALS13 | ENSG00000105198 | ECM-affliated proteins |
| LGALS14 | ENSG00000006659 | ECM-affliated proteins |
| LGALS16 | ENSG00000249861 | ECM-affliated proteins |
| LGALS2 | ENSG00000100079 | ECM-affliated proteins |
| LGALS3 | ENSG00000131981 | ECM-affliated proteins |
| LGALS4 | ENSG00000171747 | ECM-affliated proteins |
| LGALS8 | ENSG00000116977 | ECM-affliated proteins |
| LGALS9 | ENSG00000168961 | ECM-affliated proteins |
| LGALS9B | ENSG00000170298 | ECM-affliated proteins |
| LGALS9C | ENSG00000171916 | ECM-affliated proteins |
| LMAN1 | ENSG00000074695 | ECM-affliated proteins |
| LMAN1L | ENSG00000140506 | ECM-affliated proteins |
| MBL2 | ENSG00000165471 | ECM-affliated proteins |
| OVGP1 | ENSG00000085465 | ECM-affliated proteins |
| PARM1 | ENSG00000169116 | ECM-affliated proteins |
| OPRPN | ENSG00000171199 | ECM-affliated proteins |
| REG1A | ENSG00000115386 | ECM-affliated proteins |
| REG1B | ENSG00000172023 | ECM-affliated proteins |
| REG3A | ENSG00000172016 | ECM-affliated proteins |
| REG3G | ENSG00000143954 | ECM-affliated proteins |
| REG4 | ENSG00000134193 | ECM-affliated proteins |
| ITGAD | ENSG00000156886 | Integrins |
| ITGAE | ENSG00000083457 | Integrins |
| ITGAL | ENSG00000005844 | Integrins |
| ITGAV | ENSG00000138448 | Integrins |
| ITGA1 | ENSG00000213949 | Integrins |
| ITGA2 | ENSG00000164171 | Integrins |
| ITGA2B | ENSG00000005961 | Integrins |
| ITGA3 | ENSG00000005884 | Integrins |
| ITGA4 | ENSG00000115232 | Integrins |
| ITGA5 | ENSG00000161638 | Integrins |
| ITGA6 | ENSG00000091409 | Integrins |
| ITGA7 | ENSG00000135424 | Integrins |
| ITGA8 | ENSG00000077943 | Integrins |
| ITGA9 | ENSG00000144668 | Integrins |
| ITGA10 | ENSG00000143127 | Integrins |
| ITGA11 | ENSG00000137809 | Integrins |
| ITGB1 | ENSG00000150093 | Integrins |

| | | |
|---|---|---|
| ITGB3 | ENSG00000259207 | Integrins |
| ITGB4 | ENSG00000132470 | Integrins |
| ITGB5 | ENSG00000082781 | Integrins |
| ITGB6 | ENSG00000115221 | Integrins |
| ITGB7 | ENSG00000139626 | Integrins |
| ITGB8 | ENSG00000105855 | Integrins |
| PECAM1 | ENSG00000261371 | Endothelial |
| KDR | ENSG00000128052 | Endothelial |
| NRP1 | ENSG00000099250 | Endothelial |
| CD34 | ENSG00000174059 | Endothelial |
| AQP4 | ENSG00000171885 | Astrocyte |
| MLC1 | ENSG00000100427 | Astrocyte |
| GFAP | ENSG00000131095 | Astrocyte |
| ALDH1L1 | ENSG00000144908 | Astrocyte |
| HEPACAM | ENSG00000165478 | Astrocyte |
| RBFOX3 | ENSG00000167281 | Neural |
| CALB1 | ENSG00000104327 | Neural |
| CALB2 | ENSG00000172137 | Neural |
| S100G | ENSG00000169906 | OPC |
| SCGN | ENSG00000079689 | Neural |
| NEFL | ENSG00000277586 | Neural |
| NEFH | ENSG00000100285 | Neural |
| NEFM | ENSG00000104722 | Neural |
| AIF1 | ENSG00000204472 | Microglia |
| CD68 | ENSG00000129226 | Microglia |
| TMEM119 | ENSG00000183160 | Microglia |
| OLIG2 | ENSG00000205927 | OPC |
| SOX10 | ENSG00000100146 | Oligodendrocytes |
| MBP | ENSG00000197971 | Oligodendrocytes |
| VIM | ENSG00000026025 | GBM |
| EGFR | ENSG00000146648 | GBM |
| IDH1 | ENSG00000138413 | GBM |
| IDH2 | ENSG00000182054 | GBM |
| NES | ENSG00000132688 | GSC |
| SOX2 | ENSG00000181449 | GSC |
| GRIP1 | ENSG00000155974 | Neural |
| GRIP2 | ENSG00000144596 | Neural |
| NPNT | ENSG00000168743 | Neural |
| CUX2 | ENSG00000111249 | Neural |
| TESPA1 | ENSG00000135426 | Neural |

| | | |
|---|---|---|
| ZBBX | ENSG00000169064 | Neural |
| ADAMTS16 | ENSG00000145536 | Neural |
| FSTL4 | ENSG00000053108 | Neural |
| OTOGL | ENSG00000165899 | Neural |
| POU6F2 | ENSG00000106536 | Neural |
| RORB | ENSG00000198963 | Neural |
| SLC17A6 | ENSG00000091664 | Neural |
| PCSK1 | ENSG00000175426 | Neural |
| RIT2 | ENSG00000152214 | Neural |
| SNCG | ENSG00000173267 | Neural |
| SORCS1 | ENSG00000108018 | Neural |
| HTR2A | ENSG00000102468 | Neural |
| RXFP1 | ENSG00000171509 | Neural |
| ATP2C2 | ENSG00000064270 | Neural |
| CD36 | ENSG00000135218 | Neural |
| CRYM | ENSG00000103316 | Neural |
| HTR2C | ENSG00000147246 | Neural |
| TPH2 | ENSG00000139287 | Neural |
| TSHZ2 | ENSG00000182463 | Neural |
| VWC2L | ENSG00000174453 | Neural |
| FILIP1 | ENSG00000118407 | Neural |
| HS3ST2 | ENSG00000122254 | Neural |
| HS3ST4 | ENSG00000182601 | Neural |
| KCNH5 | ENSG00000140015 | Neural |
| ADAMTS3 | ENSG00000156140 | Neural |
| SNTB2 | ENSG00000168807 | Neural |
| CDH12 | ENSG00000154162 | Neural |
| GAS2L3 | ENSG00000139354 | Neural |
| NPY1R | ENSG00000164128 | Neural |
| NTNG2 | ENSG00000196358 | Neural |
| NWD2 | ENSG00000174145 | Neural |
| RASGRP1 | ENSG00000172575 | Neural |
| SLC26A4 | ENSG00000091137 | Neural |
| THEMIS | ENSG00000172673 | Neural |
| TRPC5 | ENSG00000072315 | Neural |
| ZDHHC23 | ENSG00000184307 | Neural |
| NPFFR2 | ENSG00000056291 | Neural |
| ROS1 | ENSG00000047936 | Neural |
| SLC17A7 | ENSG00000104888 | Neural |
| DNER | ENSG00000187957 | Neural |
| PTCHD4 | ENSG00000244694 | Neural |
| ADRA1A | ENSG00000120907 | Neural |
| ADRA1B | ENSG00000170214 | Neural |

| | | |
|---|---|---|
| LYPD6B | ENSG00000150556 | Neural |
| PROX1 | ENSG00000117707 | Neural |
| THSD7B | ENSG00000144229 | Neural |
| COL25A1 | ENSG00000188517 | Neural |
| PLCH1 | ENSG00000114805 | Neural |
| SYNPR | ENSG00000163630 | Neural |
| TRHDE | ENSG00000072657 | Neural |
| SLC24A3 | ENSG00000185052 | Neural |
| VIP | ENSG00000146469 | Neural |
| NLGN3 | ENSG00000196338 | Neural |
| BDNF | ENSG00000176697 | Neural |
| GRP78 | ENSG00000044574 | Neural |
| ADAM10 | ENSG00000137845 | Neural |
| IGF-1 | ENSG00000017427 | Neural |
| NTRK2 | ENSG00000148053 | Neural |
| PIK3CA | ENSG00000121879 | Neural |
| GAP43 | ENSG00000172020 | Neural |
| B4GALNT1 | ENSG00000135454 | GBM |
| BCAN | ENSG00000132692 | GBM |
| CAV1 | ENSG00000105974 | GBM |
| CHODL | ENSG00000154645 | GBM |
| ELOVL2 | ENSG00000197977 | GBM |
| HES1 | ENSG00000114315 | GBM |
| HILPDA | ENSG00000135245 | GBM |
| IGFBP3 | ENSG00000146674 | GBM |
| IGFBP5 | ENSG00000115461 | GBM |
| LOX | ENSG00000113083 | GBM |
| MGST1 | ENSG00000008394 | GBM |
| NNAT | ENSG00000053438 | GBM |
| PSEN2 | ENSG00000143801 | GBM |
| PSENEN | ENSG00000205155 | GBM |
| SOX11 | ENSG00000176887 | GBM |
| SOX4 | ENSG00000124766 | GBM |
| TP53 | ENSG00000141510 | GBM |
| TRIL | ENSG00000255690 | GBM |
| TTYH1 | ENSG00000167614 | GBM |
| APOE | ENSG00000130203 | Microglia-PVM |
| ARHGAP24 | ENSG00000138639 | Microglia-PVM |
| CCL4 | ENSG00000275302 | Microglia-PVM |
| CD86 | ENSG00000114013 | Microglia-PVM |

| | | |
|---|---|---|
| CTSH | ENSG00000103811 | Microglia-PVM |
| FASLG | ENSG00000117560 | Microglia-PVM |
| HLA-DQA1 | ENSG00000196735 | Microglia-PVM |
| IFITM3 | ENSG00000142089 | Microglia-PVM |
| ITGAM | ENSG00000169896 | Microglia-PVM |
| ITGAX | ENSG00000140678 | Microglia-PVM |
| LRRK1 | ENSG00000154237 | Microglia-PVM |
| LYVE1 | ENSG00000133800 | Microglia-PVM |
| P2RY12 | ENSG00000169313 | Microglia-PVM |
| P2RY13 | ENSG00000181631 | Microglia-PVM |
| PTPRC | ENSG00000081237 | Microglia-PVM |
| RGS16 | ENSG00000143333 | Microglia-PVM |
| SPI1 | ENSG00000066336 | Microglia-PVM |
| TGFB1 | ENSG00000105329 | Microglia-PVM |
| TREM2 | ENSG00000095970 | Microglia-PVM |
| MKI67 | ENSG00000148773 | Proliferation |
| PCNA | ENSG00000132646 | Proliferation |
| CCNA1 | ENSG00000133101 | Proliferation |
| CCNB2 | ENSG00000157456 | Proliferation |
| CDK1 | ENSG00000170312 | Proliferation |
| CENPF | ENSG00000117724 | Proliferation |
| KIT | ENSG00000157404 | Proliferation |
| TOP2A | ENSG00000131747 | Proliferation |
